# Near-infrared MINFLUX imaging enabled by suppression of fluorophore blinking

**DOI:** 10.1101/2024.08.27.609859

**Authors:** C Venugopal Srambickal, H Esmaeeli, J Piguet, L Reinkensmeier, R Siegmund, M Bates, A Egner, J Widengren

## Abstract

MINimal photon FLUXes (MINFLUX) offers super-resolution microscopy (SRM) with nanometer localization precision, with more relaxed fluorophore brightness and photostability requirements than for other SRM techniques. Nonetheless, low localization probabilities have been reported in several MINFLUX studies, and a broader use of less bright and photostable fluorophores, including near-infrared (NIR) fluorophores has been difficult to realize. In this work, we identified fluorophore blinking as a main cause of erroneous (and dismissed) fluorophore localizations in MINFLUX imaging and devised strategies to overcome these effects. We systematically studied the blinking/switching properties of cyanine fluorophores emitting in the far-red or NIR range, and over typical time scales (µs-10ms), sample and excitation conditions used in MINFLUX imaging. By subsequent simulations of representative MINFLUX localization procedures, we found that trans-cis isomerization, and in particular photo-reduction of the fluorophores, can generate significant localization errors. However, these localization errors could be suppressed by balanced redox buffers and repetitive excitation beam scans. Implementing these strategies, and replacing the slower, intrinsic switching of the fluorophores needed for the localization by transient binding of fluorophore-labelled DNA strands to complementary DNA strands attached to the targets (DNA-PAINT), we could for the first time demonstrate NIR-MINFLUX imaging with nanometer localization precision. This work presents an overall strategy, where fluorophore blinking characterization and subsequent simulations make it possible to design optimal sample and excitation conditions, opening for NIR-MINFLUX imaging, as well as for a broader use of fluorophores in MINFLUX and related SRM studies.

## Introduction

Following the development of far-field super-resolution microscopy (SRM), including coordinate-targeted stimulated emission depletion (STED)^1, 2^ and single molecule localization microscopy (SMLM),^3, 4^ MINimal photon FLUXes (MINFLUX)^5, 6^ represents a next generation SRM technique taking fluorophore localization precision to new levels, with less fluorescence photons required for the localization. The more relaxed requirements on molecular brightness and photostability in MINFLUX can open for the use of new categories of fluorophores, in particular near-infrared (NIR) fluorophores, which due to limited photon budgets have found a quite minor use in SRM this far. In MINFLUX, the fluorophores to be localized are first switched individually, typically on a slower time scale of milliseconds or longer, like in SMLM. However, while in SMLM the localizations of individual, spatially separable, emitting fluorophores are inferred from the centroids of their emitted fluorescence as imaged by a camera, the MINFLUX concept uses a movable excitation beam with an intensity minimum, e.g. a donut beam as used in STED imaging. The coordinates of individual fluorophores can then be inferred from the detected fluorescence photon numbers relative to one another at the different beam positions, forming a so-called targeted coordinate pattern (TCP) around the fluorophore to be localized. The excitation intensity experienced by a fluorophore will be weaker the closer it is to the minimum of the excitation beam for a laser beam position within a TCP. Ideally, if the excitation minimum is zero, the absence of fluorescence emission can provide a signature that the excitation zero coincides spatially with the fluorophore. MINFLUX, acknowledged as the most efficient known way of using fluorescence photons for localizations,^7^ thus opens for the use of emitters with lower photon budgets (lower molecular brightness and photostability), but on the other hand also puts other requirements in the forefront, such as low background levels.^5^

In recent years, fluorescence imaging in the near-infrared (NIR) range has received increased attention and comes with several advantages, including reduced scattering, lower absorption, autofluorescence and phototoxicity in the sample, as well as deeper penetration depths.^8-10^ Given the relatively strict requirements on fluorophore photostability and brightness in SRM,^11,12^ and since organic fluorophores generally display lower fluorescence brightness and photostability the more red-shifted their excitation and emission spectra, NIR fluorophores have hitherto found limited use in SRM. However, in MINFLUX imaging the generally lower photon budgets of these fluorophores should be better tolerated. Moreover, excitation and emission in the NIR provide prerequisites for reaching lower background levels and offer additional spectral windows for simultaneous SRM imaging over multiple, spectrally separated color channels.

In the red spectral range, pentamethine cyanines such as AlexaFluor 647 (AF647) have become fluorophores-of-choice for SMLM, in particular for stochastic optical reconstruction microscopy (STORM),^4, 13^ combining the necessary fluorophore brightness and photostability with appropriate slow switching properties (number of switching cycles, on-off duty cycles).^14^ These properties and its usefulness for STORM imaging has also qualified AF647 as one of the first fluorophores to be established for MINFLUX imaging.^5, 6^ In the NIR, heptamethine cyanines represent the by far most commonly used fluorophore category for bioimaging and can be expected to show similar slow switching properties as AF647 and other pentamethine cyanines.^15^ While they typically lack the brightness and photostability to make them attractive for STORM imaging, it can be argued that they may still qualify for MINFLUX imaging, for which the photon budget requirements are lower. However, even in the visible range, using fluorophores with better photon budgets, it has been noted that current MINFLUX implementations tend to detect fewer fluorophores, i.e. to show lower probabilities of localizing fluorophore-labeled target molecules, compared to camera-based (widefield) SMLM techniques.^7, 16^ It is unlikely that these lowered probabilities are related to the MINFLUX concept itself,^17^ but rather to properties of the fluorophores used. In the MINFLUX localization procedure, the fluorophore to be localized must first switch in a slow (milliseconds-seconds) time scale, and with a duty cycle low enough (typically 10^−3^-10^−4^) to allow individual, well separated fluorophores to be targeted in the localization. This requirement is shared with SMLM^14^ and should thus not lead to a lowered localization probability in MINFLUX compared to in SMLM. In the next MINFLUX localization step(s), however, with a donut-shaped excitation beam scanned around the emissive fluorophore in a pre-defined TCP, it is typically assumed that the fluorescence signal is linearly proportional to the excitation intensity experienced by the fluorophore. If the fluorophore would undergo faster blinking/switching between different scanning positions of the donut beam, however, this linearity assumption is no longer fully valid, and the ability to infer the fluorophore localization from its emitted fluorescence at each scanning position may be compromised. Such effects can in fact be a major reason for the lowered localization probabilities experienced in MINFLUX imaging, as recently reported.^7, 16^ Accordingly, given the similarities in blinking/switching properties between NIR heptamethine and far-red pentamethine cyanines such as AF647, blinking/switching may also be a major complication for the use of heptamethine cyanines, and possibly a major hurdle needed to overcome to make NIR-MINFLUX imaging possible. While effects of fluorophore blinking in MINFLUX measurements has been brought up as a possible complication,^18^ a more comprehensive investigation of such effects has to our knowledge not been reported. Such investigation is relevant for MINFLUX imaging, but also to other SRM techniques incorporating time-modulated illumination patterns in the localization,^19-21^ and can provide an understanding, whereby imaging strategies can be modified to minimize or eliminate localization errors. Consequently, in this work, we performed systematic studies of far-red and NIR cyanine fluorophore blinking with subsequent simulations of MINFLUX localizations. This allowed us to design optimal sample and excitation conditions and then to demonstrate NIR-MINFLUX imaging in practice. More generally, this work can thereby also open for a broader use of NIR fluorophores in MINFLUX and related SRM studies.

## Results and Discussion

To understand how and to what extent fluorophore blinking can affect localization precisions and probabilities in MINFLUX imaging, and to develop possible remedy strategies, we systematically studied the blinking/switching properties of the pentamethine cyanine AF647, over a time scale spanning several orders of magnitude (µs-10ms), and under labeling, sample and excitation conditions relevant for MINFLUX imaging. Similarly, we also studied three heptamethine cyanine fluorophores; Dylight 755 (DL755), AlexaFluor 750 (AF750) and CF750 (CF750), as possible candidate fluorophores for MINFLUX imaging in the NIR, displaying similar (STORM compatible) slow switching properties as AF647.^14^ From the acquired data, photodynamic models with rate parameters were determined, based on which we then performed simulations of the fluorophore photodynamics and blinking behavior, under representative MINFLUX excitation beam scans and sample conditions. From these simulations, we could identify major experimental conditions and fluorophore blinking properties compromising the localization precision and probability, allowing us to formulate strategies to minimize these effects. Finally, we then applied these strategies in practice, demonstrating NIR MINFLUX imaging for the first time, with nanometer localization precision.

### Photophysical characterization

We applied transient state (TRAST) spectroscopy, offering a robust approach to monitor fluorophore blinking kinetics over a µs to 10ms time range,^22, 23^ performed on a customized wide-field microscope setup, with laser excitation at 640nm (for AF647) or 750nm (for DL755, AF750 and CF750) (See Methods). By so-called TRAST curves, plotting the time-averaged fluorescence intensity, ⟨*F*_*exc*_(*w*)⟩_*norm*_, generated by rectangular excitation pulse trains with different pulse widths, *w*, characteristic relaxations on a μs to ms time scale were recorded in the fluorophore samples. These relaxations reflect build-up of photoinduced non- or weakly emissive states in the fluorophores, generated following the onset of a rectangular excitation pulse (see SI Section S1, Eqs. S1-S4). From the relaxations recorded in the TRAST curves, we established a model for the underlying state transitions in the fluorophores and determined their transition rates under different excitation and sample conditions relevant for MINFLUX imaging and SMLM.

To assign a photophysical model, we first performed TRAST experiments on free fluorophores in an air-saturated aqueous buffer solution (11.8 mM PBS, pH 7.4). TRAST curves recorded from AF647 and the three NIR cyanines all displayed a major relaxation process in the faster µs-100µs time range, as well as a prominent slower (1-10ms) relaxation (Figure 1A). When applying pulse trains with different excitation photon fluxes, Φ_*exc*_, the amplitude of the faster (µs-100µs) relaxation remained largely constant (between 25-40% for the different fluorophores), while its relaxation time decreased with higher Φ_*exc*_ (Figures 1B and 1C, SI section S2, Figure S1). This Φ_*exc*_-dependence is consistent with reversible excitation-driven isomerization, as previously observed in fluorescence correlation spectroscopy (FCS)^24, 25^ and TRAST experiments.^26, 27^ Since isomerization can take place over several bonds in the polymethine chains of the fluorophores, different photo-isomerized states can likely be populated.^26^ However, compared to isomerization to and from an emissive all-*trans* state, N, transitions between different (non- or weakly emitting) photo-isomerized states yield minor blinking effects. A single state representing all photo-isomerized states, P, is thus sufficient to describe the blinking kinetics, as observed in the TRAST experiments, and as a basis for the simulations of the MINFLUX localization. As seen in the TRAST experiments, and since the N-P transitions are excitation-driven, P can get prominently (25-40%) populated over a broad range of excitaiton intensities. For the slower (1–10ms) relaxation, we observed an increase in its amplitude and a faster decay with higher Φ_*exc*_ (Figures 1B and 1C). For many cyanine fluorophores photo-reduction is a major dark state transition, reported to take place from the triplet state,^28, 29^ and the Φ_*exc*_-dependence observed in the TRAST curves for this relaxation is well in agreement with such transition, with a dark state recovery which is not excitation-driven. In the TRAST curves (Figures 1B and 1C), no significant build-up of the triplet state itself was observed, also not at higher Φ_*exc*_. This is consistent with previous FCS and TRAST studies of penta- and heptamethine cyanines,^24, 26, 27^ and indicates that intersystem crossing is outcompeted by the higher photo-isomerization rates in these fluorophores. Low triplet state populations are also expected, given the relatively short excited state lifetimes of the fluorophores (1.0 ns for AF647, ∼0.5 ns for the NIR cyanines, see SI Section S3), resulting in low excited singlet state populations for the Φ_*exc*_ applied in our TRAST experiments, and small effective rates of intersystem crossing compared to the triplet state decay rates in presence of molecular oxygen (typically ∼0.5µs^-1^ in air-saturated aqueous solutions).^24, 30^

**Figure 1:**
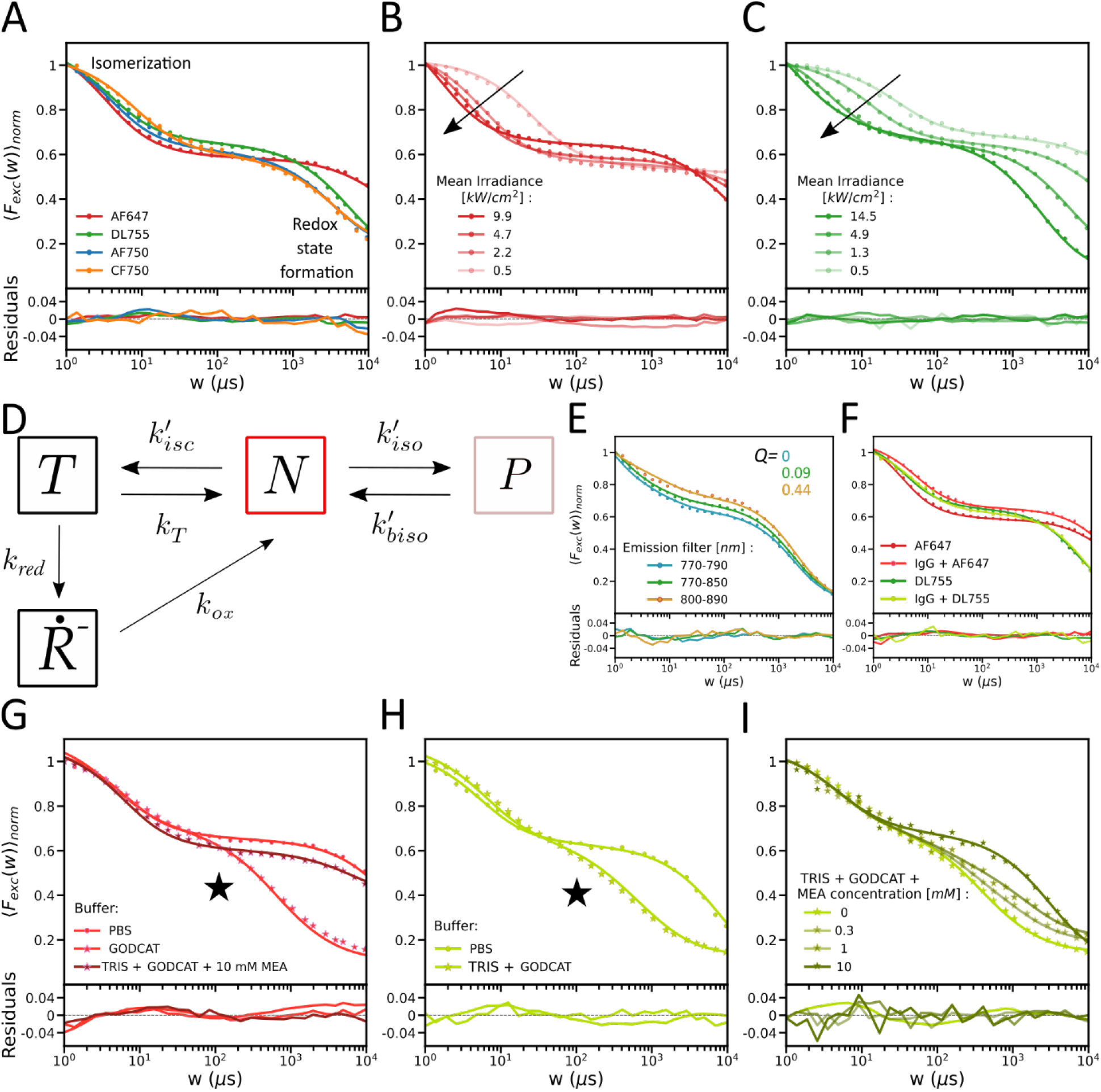
Recorded TRAST curves, with experimental data represented by dots, and fitted curves with lines. Fitting residuals are plotted below. If not stated otherwise, the TRAST curves were recorded in air-saturated PBS solution, with excitation at 750nm for the NIR dyes and at 638nm for AF647. A) TRAST curves recorded from all NIR dyes (ϕ_exc_= 4.9kW/cm^2^) and from AF647 (ϕ_exc_= 4.7kW/cm^2^). Two major relaxations are observed, attributed to photo-isomerization and redox state formation. B) Excitation intensity dependence for AF647. Arrow indicates higher intensities. C) Excitation intensity dependence for DL755. Arrow indicates higher intensities. D) Photophysical model, as described in the main text and used to fit the experimental TRAST curves. E) TRAST curves recorded from DL755 with different emission filters (ϕ_exc_= 14.5kW/cm^2^). F) TRAST curves recorded from AF647(ϕ_exc_= 4.7kW/cm^2^) and DL755(ϕ_exc_= 4.9kW/cm)^2^, in free form and when conjugated to an antibody (IgG). G) TRAST curves recorded from AF647, in PBS, when in a deoxygenated buffer (Tris+GODCAT) and upon addition of MEA (10mM), (ϕ_exc_= 4.7kW/cm^2^). Black star: triplet state buildup. H) Effects of deoxygenation (GODCAT) in DL755, (ϕ_exc_= 4.9kW/cm^2^). Black star: triplet state buildup. I) TRAST curves recorded from DL755 in GODCAT with different concentrations of MEA added (ϕ_exc_-4.9kW/cm^2^).

At the time scale of the fluorophore dark state transitions and the TRAST experiments (1µs-10ms), and at which time scale also the MINFLUX beam localization procedure typically operates, ground and excited singlet state relaxations (ns) within N and P will be averaged out. For the fluorophores studied, we could thus assign a photophysical model, including an all-*trans* (N), photo-isomerized (P), triplet (T) and photo-reduced 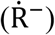 state (Figure 1D) to describe the observed relaxations in the TRAST curves (Figures 1A-C), and a fluorophore state vector 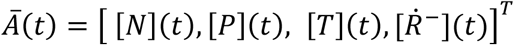 representing the population probabilities of the N, P, T and 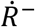 states (SI Section S4). In recorded TRAST curves from DL755, we found that the amplitude of the fast, photo-isomerization relaxation was lowered the more red-shifted emission band-pass filters were used (Figure 1E). We attribute this amplitude effect to weak, red-shifted emission from photo-isomerized states of the fluorophores, as recently identified for other cyanine fluorophores.^26, 27^ It can be accounted for in the photophysical model by assigning P a relative brightness factor, *Q* (Eq. S13). In absence of excitation, polymethine cyanine fluorophores typically exist in their all-*trans* state.^29^ From our model (Figure 1D), with fluorophores subject to constant Φ_*exc*_ starting at time *t*=0 we can then set 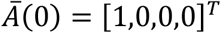 as initial condition. Relaxations in the generated fluorescence intensity, *F*(*t*), as observed in TRAST curves (see Section S5, Eqs. S14-S16), can then be attributed to how [*N*] and [*P*] evolves within rectangular excitation pulses with increasingly longer pulse durations:

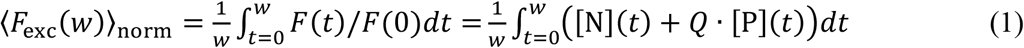

To further evaluate the model of Figure 1D, fitting of photophysical rate parameters was performed as previously described.^26, 27, 31-33^ Herein, theoretical TRAST curves were calculated based on Eq. 1, and with the ⟨*F*_*exc*_(*w*)⟩_*norm*_ data points of the TRAST curves calculated from the recorded wide-field data as described in SI, Section S5. The set of rate parameter values best reproducing the experimental data was then found using nonlinear least-squares optimization (SI, Section S6). For each of the fluorophore samples, global fitting of rate parameters to multiple TRAST curves, recorded at different Φ_*exc*_ and with different emission filters could well reproduce the experimental data (Figures 1B, 1C and 1E and SI Section S2). The fitted parameters values are listed in Section S7, Table S1.

Next, we studied how the photophysical transitions were affected by the sample conditions used for STORM and MINFLUX imaging. Compared to free fluorophores, conjugation to antibodies resulted in longer isomerization relaxation times (Figure 1F, Figure S3 in Section S8) reflecting lower isomerization and back-isomerization rates (Table S1). This agrees with previous observations that bulky substituents on the cyanine head groups can increase the viscous drag, retarding conformational reorganizations within the molecule as well as rates for photo-induced isomerization and back-isomerization.^24, 29^ We then investigated effects of STORM imaging buffers on the fluorophore blinking kinetics of the antibody-conjugated fluorophores. Such buffers are designed to reversibly transit fluorophores in the illumination field to dark states, thereby restricting the fraction of simultaneously active fluorophores while at the same time maintaining a high brightness in the active state.^34^ For AF647 and other cyanines, these buffers typically contain an enzyme-based oxygen scavenger system (GODCAT, see Methods), together with a thiol-based reductant, such as mercaptoethylamine (MEA). The recorded TRAST curves largely display the expected effects of these imaging buffer constituents; Deoxygenation by GODCAT leads to a prominent increase in the triplet state (T) buildup, taking place at an order of magnitude faster time scale than photo-reduction (Figures 1G and 1H, black star). Titration of MEA into the deoxygenated buffer resulted in depopulation of T into the 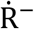 state (Figure 1G and 1I, SI Section S8, Fig S3). Notably, in the recorded TRAST curves (Figures 1G and 1H), the 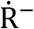 relaxation occurs on a much (more than an order of magnitude) faster time scale than for the long lived dark state in STORM experiments.^14^ For cyanines in their triplet state, thiols can promote photo-induced electron transfer, with subsequent triplet-to-singlet intersystem crossing in geminate cyanine-thiol radical pairs. Then, either back-electron transfer into emissive singlet state fluorophores can take place, or more long-lived, non-emissive cyanine-thiol adducts are formed.^35^ The latter forms the basis for the slow switching required for STORM measurements,^13^ while our interpretation is that the TRAST measurements mainly shows the accompanying photo-reduction attributed to collisional encounters with MEA, with no subsequent adduct formation and a much more short-lived, non-adducted 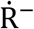 state. The experimental TRAST curves recorded from samples with GODCAT and MEA could be well reproduced by rate parameter fitting, using the same photophysical model as for the free fluorophores (Figure 1D). In the fitting, the number of freely fitted parameters were limited by fitting some rate parameters globally for multiple TRAST curves. The fitted rate parameters (Table S1) clearly reflect the effects of MEA on the blinking kinetics (Figure 1I), promoting triplet state deactivation in the GODCAT samples by reduction back to the ground singlet state (*k*_*T*_), and by an enhanced transition rate (*k*_*red*_) to a photo-reduced state.

Overall, from the TRAST experiments we find that dark or dim states are generated to a significant extent for all fluorophores, by photo-isomerization and by photo-reduction. The time scales of these transitions depend on the excitation and sample conditions but fall within typical time scales of MINFLUX localization procedures and their TCP iterations (typically tens of microseconds up to milliseconds to complete a TCP). It can thus be expected that situations can occur in MINFLUX experiments, where these transitions can perturb the linear relationship assumed between the local Φ_*exc*_ experienced by a fluorophore and its fluorescence photon emission rate.

### Simulations

To assess the effects of fluorophore blinking in MINFLUX localizations more in detail, we performed simulations with iterative TCP scans around a fluorophore. The fluorophore was simulated undergoing blinking transitions as determined from our TRAST experiments, within a basic, representative localization algorithm used in MINFLUX imaging ^6, 36^ (Figure 2A, further details in SI Section S9); A donut shaped excitation beam was translated within a two-dimensional plane in a hexagonal TCP around the fluorophore located at position, 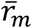(Figure 2A, gold star). In each iteration, the donut beam center (Figure 2A, red dots) was first parked in the center of the hexagonal TCP and then at each of its 6 vertices, with the same beam dwell time, *t*_*dwell*_, at each of the positions. Within each iteration (represented by rows in figure 2A), fluorescence photon counts, 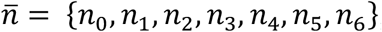, generated from the fluorophore by the excitation beam in its seven beam positions relative to 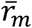 were calculated as an integral over *t*_*dwell*_ for each beam position of the TCP. The calculations were based on the photophysical model of Figure 1D, with transition rates as determined for the fluorophore (Table S1), with the all-*trans* state (N) as the only emissive state, and based on its corresponding excitation cross section, *σ*_*N*_, and fluorescence lifetime (SI Section S3). To mimic how the location of the fluorophore then is estimated we applied a maximum likelihood estimator (MLE) algorithm, as applied in MINFLUX,^5^ which estimates the fluorophore position, 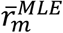, best in agreement with 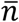. The TCP center is then relocated to the fluorophore position found by the MLE algorithm 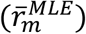 and the localization procedure is then repeated in 3 more iterations, with the diameter, *L*, of the TCP pattern reduced in each iteration step, and the excitation power increased (illustrated in the three lower rows of figure 2A, see also SI Section S9 for further details). In the simulations, the actual purpose was to analyse how blinking kinetics in the fluorophores effects the localization. We therefore did not include background contributions from other fluorophores in the sample, and also not detector losses or any collection efficiency differences. To simulate the evolution of photophysical states on a single fluorophore level, as in actual MINFLUX imaging experiments, we applied a Markovian chain model (SI, Section S9), based on the model in figure 1D and the transition rates as determined in the TRAST experiments (Table S1). Further, since it is not possible to include all possible scenarios for the localization procedure in the simulations, we confined the simulations to iterations after pre-localization, with the fluorophore location, 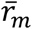, inside the TCP, and with a few representative starting excitation powers, fluorophore locations and beam dimensions considered. This allowed us to capture major effects and features following from fluorophore blinking within a limited number of representative scenarios.

**Figure 2:**
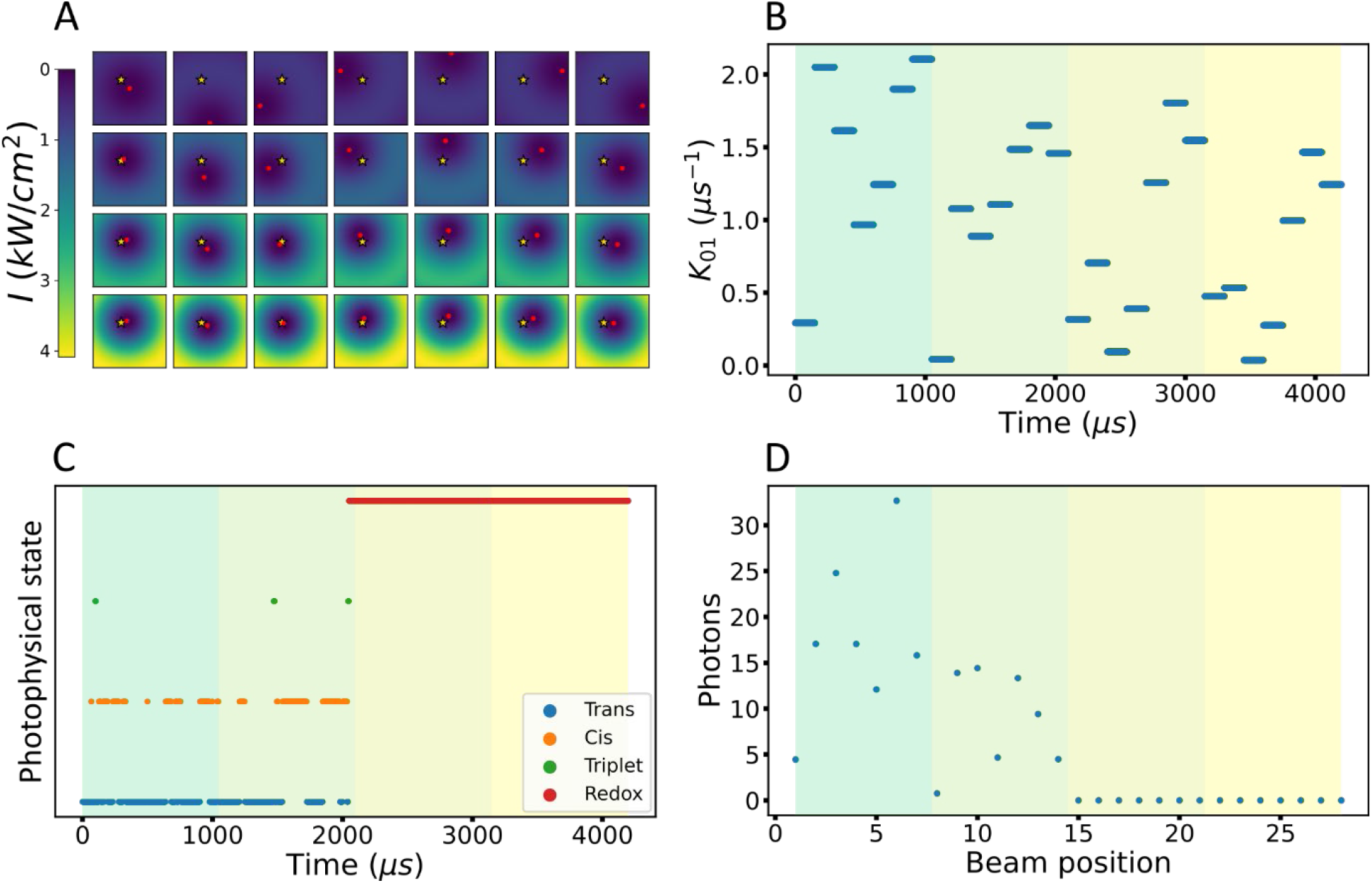
Simulation of a representative MINFLUX localization. A) Illustration of the fluorophore position (golden star, in this example located at (−35nm,35nm) with respect to the TCP center in the first iteration) and the different beam positions (beam centers marked red dots) over the different TCP iterations (corresponding to each row), with the excitation intensity increasing and the diameter of the TCP (L) decreasing from one TCP iteration to the next. B) The excitation rate (k_01_) experienced at the fluorophore position over time as the beams are moved in the four TCP iterations. C) Photophysical state evolution showing the fluorophore undergoing transitions between the bright trans isomer state (N, blue) and the dark cis isomer (P, orange), triplet (T, green) and the dark long-lived redox (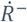, red) state during the time of localization. D) The integrated photons collected from the fluorophore for each of the beam positions shown in figure A. The different coloured regions in figure B, C and D indicate the different TCP iterations.

Given an iterative MINFLUX localization procedure applied on a single fluorophore located at 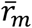 from the center of the first TCP, as shown in Figure 2A, we simulated possible photophysical state evolution outcomes for the fluorophore (DL755, labelled to an antibody and in a Tris + GODCAT buffer with 10mM MEA). A representative outcome of such simulations is shown in Figure 2. Excitation rates, as experienced by the fluorophore are calculated throughout the procedure, from the irradiance at 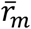 for the different beam positions within each TCP iteration (Figure 2B). From the state evolution (Figure 2C), the fluorescence photon counts, 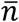, are then calculated in the simulations after each TCP iteration and for each beam position (Figure 2D). In the outcome shown in Figure 2C, the simulated fluorophore switches reversibly between its emissive, all-*trans* (N) and dark *cis* (P) state during the first two TCP iterations, and only occasionally goes into its triplet state (T). Towards the end of the second iteration, at around 2 ms, the fluorophore goes into its more long-lived redox state 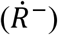 from which the fluorophore does not relax back within the simulated time. The simulated state evolution for this single fluorophore largely reflects the transient state dynamics, as observed in the TRAST experiments on an ensemble level; Upon excitation, an equilibration between N and P takes place on a 1-100µs time range, with minor triplet state buildup, followed by 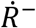 buildup on a ms time scale. In the simulated example, it can be noted that the calculated 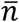 after each TCP iteration (Figure 2D) is far from linearly proportional to the corresponding excitation rates experienced by the fluorophore in each of the beam positions within the same TCP iteration (Figure 2B). This illustrates that fluorophore blinking properties of relevant fluorophores and under relevant experimental conditions for MINFLUX imaging indeed can distort the calculation of 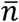. This distortion will affect the MLE estimation of 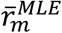 for the next iteration, and errors due to fluorophore blinking may thus propagate and accumulate in the localization procedure. Given that the fluorophore blinking kinetics depend on the local Φ_*exc*_ experienced by the fluorophore and the duration of the excitation, several parameters can influence the localization in MINFLUX experiments, including 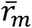 (with respect to the TCP), *t*_*dwell*_, laser power(s) applied in the TCP iterations, and what TCPs (and diameters *L*) are used. When averaged over many single molecules, on an ensemble level, the population kinetics of the photophysical states will be reproducible for a given set of parameters (as verified also for our single fluorophore simulations, see SI Section 10, Figures S5 and S6). However, due to the stochastic nature of single molecules, the blinking kinetics and state evolution will vary from one individual fluorophore to another, even if subject to the same experimental conditions.

To further assess how these parameters can affect the localization, we simulated 1000 localization outcomes for the same fluorophore (DL755) as simulated in Figure 2, located at position (1nm,1nm) near to the TCP center (0,0) at the start of the first iteration (Figure 3). This round of simulations was repeated for different beam dwell times, *t*_*dwell*_, and starting laser powers, including settings typically used in experimental realizations. Figure 3A shows the resulting spread for the 1000 simulated localization estimates (the 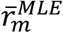 estimated by the MLE algorithm after the last TCP iteration) with respect to the actual position 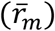, and for the different settings. In addition, we also considered localization algorithms based on multiple pattern repeats within a TCP iteration, wherein fluorescence photon counts from each beam position, 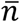, are accumulated over the repeats, using the same MLE localization procedure, beam dwell times, and starting powers as in Figure 3A, but with the dwell times distributed over 5 pattern repeats (*t*_*dwell*_ divided by 5 in each repeat) (Figure 3B, dashed boxes). Overall, the simulation outcomes (Figures 3A and 3B) suggest that fluorophore blinking can significantly affect the localization precision in MINFLUX experiments. Depending on the conditions, different dark state transitions contribute to a larger or lesser extent to the localization error. For the shorter *t*_*dwell*_ (5µs or 30µs) and lower powers (1-10µW, corresponding to peak irradiances of 0.7 – 6.8 kW/cm^2^) (red-marked squares in figure 3A), the total possible excitation time after a completed localization procedure, *t*_*total*_, will typically be smaller than the time needed for 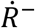 buildup in the fluorophore (four TCP iterations, each with seven beam positions correspond to a total possible beam exposure time of *t*_*total*_ = 4 × 7 × *t*_*dwell*_). The major part of the spread in the 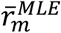 distributions is thus rather caused by photo-isomerization, between N and P, which takes place on a time scale well within *t*_*total*_. The localization precision under these conditions improved with the number of TCP iterations (SI Section S11, Figure S7). Similarly, comparing localization procedures with short *t*_*dwell*_ and low powers between Figure 3A and Figure 3B (red boxes), additional improvement in precision can be seen when N and P populations are averaged over several (here five) pattern repeats. This is consistent with observations that by spreading the beam dwell time, *t*_*dwell*_, at each beam position over multiple repeats one may obtain an averaging effect of the blinking kinetics.^18^ Next, for localization procedures with longer *t*_*dwell*_ (150-500µs), and especially when combined with higher excitation powers applied (10-30µW), an additional crescent-shaped localization error pattern emerges, displaying an appreciable loss of accuracy in the localizations (Figures 3A and 3B). Within the longer *t*_*dwell*_ applied, multiple transitions between N and P can take place and efffects on 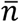 from photo-isomerization will be largely averaged out. However, the much slower transitions to and from 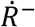 will not average out to the same extent. Yet, relaxation into 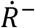 can be expected to take place well before the localization procedure is completed (*t*_*total*_). In the simulations shown in Figures 3A and 3B, the fluorophore is positioned very close (1nm,1nm) to the TCP center (0,0) at iteration 1. Nonetheless, large localization inaccuracies can still be generated, likely from buildup of the long-lived 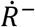 state. This will skew 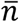 and will thereby move the TCP center to a next estimated position far from the actual fluorophore position. Once generated, the long-lived 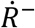 state may often not relax back within *t*_*total*_. Then, 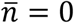 and subsequent iterations cannot correct the estimated fluorophore position (SI Section S11, figure S8). Implementing the pattern repeat leads to an overall improvement in the localization, with less inaccurate localizations, and (except for the 500µs/30µW scenario) most localizations centered around the actual fluorophore position (Figure 3B compared to Figure 3A). This improvement is likely not a result of an overall lowered 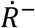 population generated, but rather suggests that with pattern repeats there is time to detect fluorescence over a larger number of beam positions before relaxation into the 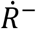 state takes place. Thereby, the skewing effect due to (unsymmetric) 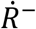 populations in the fluorophore within a TCP interation will be reduced. However, also with the pattern repeat applied, a considerable fraction of the localizations remained inaccurate. From the localization error patterns of DL755 (Figure 3A and 3B), using the lowest *t*_*dwell*_ (5µs) and laser power (1µW) seems to minimize the localization errors. However, in practice, very few fluorescence photons would be generated with such settings, which would make it difficult to distinguish these photons above typical background levels.

**Figure 3:**
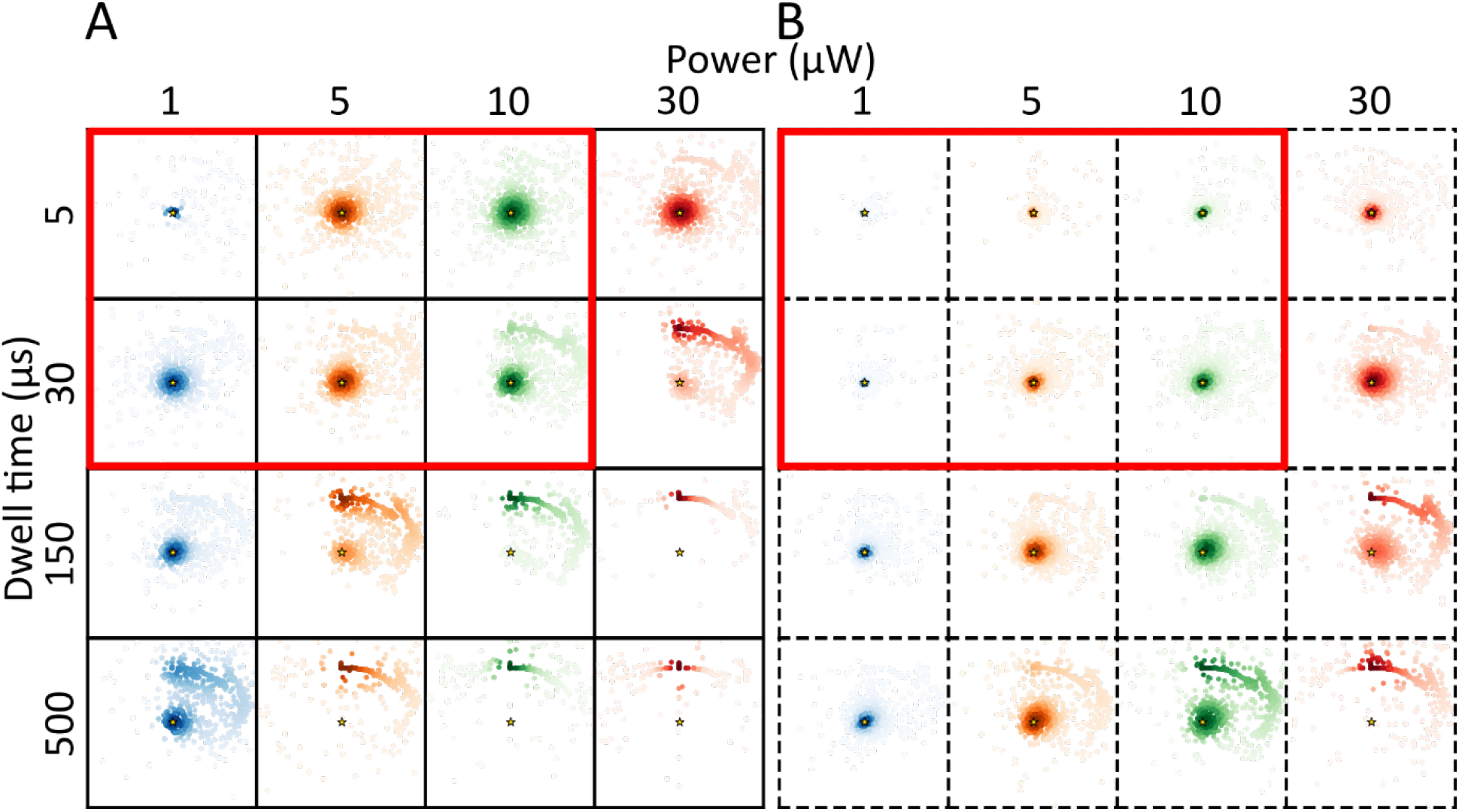
Maps of simulated final estimated locations (coloured dots) for DL755 fluorophore at position (1,1) (golden star) following an iterative MINFLUX localization. The figure shows simulated localization estimates for different excitation powers, different beam dwell times (times spend at each of the beam positions within the TCP in each of the iterations) and without (A) and with (B) pattern repeats (dotted boxes, pattern repeat of 5). The ROI (each box) has the dimensions -100nm to +100nm in the x and y directions. The difference in color transparency shows the density of estimated locations, calculated using a kernel density estimate (KDE) with a Gaussian kernel.

For reference, we also performed the same set of simulations for AF647, for the same conditions, and based on the photophysical transition rates, as determined for AF647 (Table S1). The resulting simulated localizations are displayed in SI Section S12, figures S9A and S9B. While we find similar overall features in the localization error patterns as for DL755, there are also differences attributed to the different blinking properties of the fluorophores (Table S1), which may also suggest different settings to be used in the localization procedures. For the shorter *t*_*dwell*_ and lower excitation powers applied, for which photo-isomerization is a major reason for the localization errors in DL755, AF647 displays somewhat larger errors than DL755. This likely reflects the faster and more prominent buildup of photo-isomerized P states in AF647 (Figure 1G versus Figure 1I)). On the other hand, for longer *t*_*dwell*_ and higher excitation powers, under which 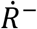 buildup is a major source of error in DL755, a lower fraction of inaccurate localizations was found in the AF647 simulations. This can likely be attributed to the comparatively lower extent of 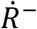 state formation in AF647 compared to in DL755 (see Figures 1G and 1I). For AF647, we find that the pattern repeat improved the localizations more than for DL755 (Figure S9B), consistent with its more prominent photoisomerization and lower tendency to undergo photoreduction (which would be less remedied by the pattern repeat).

In commercial MINFLUX instruments, automatized, FPGA-based control and assessment of the photon statistics of a localized fluorophore in between iterations are typically implemented. ^36^ By such algorithms, inaccurate localizations as found in the DL755 and AF647 simulations might then prompt the system to abort further TCP iterations and look for new fluorophores available in the field of view. This could be a possible reason for the lower final localizations observed in some MINFLUX experiments.^7, 16^ Notably, the spatial distribution of these inaccurate localizations around the correct fluorophore location is largely influenced by the excitation history, at what stage in a TCP iteration the fluorophore transits into a dark 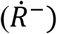 state, and the beam positions preceding that stage. To illustrate this, we performed simulations in which the TCP direction was reversed, compared to the simulations in Figure 3 (SI Section S13, Figure S10). Similarly, the actual position of the fluorophore with respect to the center of the TCP at iteration 1 will also influence the excitation history and then affect how inaccurate localizations distribute in space around the fluorophore location (SI Section S13, Figure S11).

It is worth noting that localization errors attributed to fluorophore blinking, as simulated above, are closely linked to the single fluorophore readout practiced in MINFLUX localization procedures, which places the fluorophore in a photophysical non-equilibrium. If we would consider an ensemble of DL755 fluorophores to occupy the position (1nm,1nm) instead of a single fluorophore, the photophysical states will largely reach ensemble equilibria as the localization and TCP iterations progress. The fluorescence generated from the fluorophore ensemble will then show a closer to linear dependence on the excitation rate experienced at its location (SI Section S14, Figure S12). Assuming fluorophore ensemble instead of single molecule behaviour, results in more accurate and precise localizations (SI Section S14, figure S13), in some instances even similar to that of a non-blinking single fluorophore (SI Section S14, figure S14). Bringing in ensemble features by multi-fluorophore labelling of molecular targets, or by embedding multiple fluorophores into beads would thus in principle offer a strategy to reduce localization errors in MINFLUX imaging. However, while the fluorophores should undergo blinking, they need to blink independently and then likely cannot be too close to one another. At the same time, the more separated the fluorophores, the more extended is their (common) target location, which also would compromise localization precision.

### Strategies to overcome blinking-induced localization errors

The simulations of the localization procedures above suggest that fluorophore blinking can significantly contribute to localization errors in MINFLUX imaging experiments, and for the studied cyanine fluorophores, that transitions to 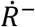 seems to be the major reason for such blinking-induced localization errors. To further assess this, we simulated 1000 localizations of a hypothetical DL755, for the same conditions as for the simulations above, with isomerization kinetics as determined from the TRAST measurements (Figure 1I, Table S1), but with no transitions to T or 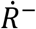. For this scenario, we find that the localization precision is better for all the conditions tested, and with negligible inaccurate localizations (Figure 4A). The localization precision improved with longer *t*_*dwell*_, higher laser power, and with the pattern repeat included (Figure 4B). This is consistent with a larger extent of averaging and/or equilibration of the P buildup of the DL755 that follows for each beam location within a TCP, with an increase in beam dwell time, with higher excitation intensities (and shorter isomerization relaxation times), as well as when this buildup is averaged over a TCP iteration by several pattern repeats.

**Figure 4:**
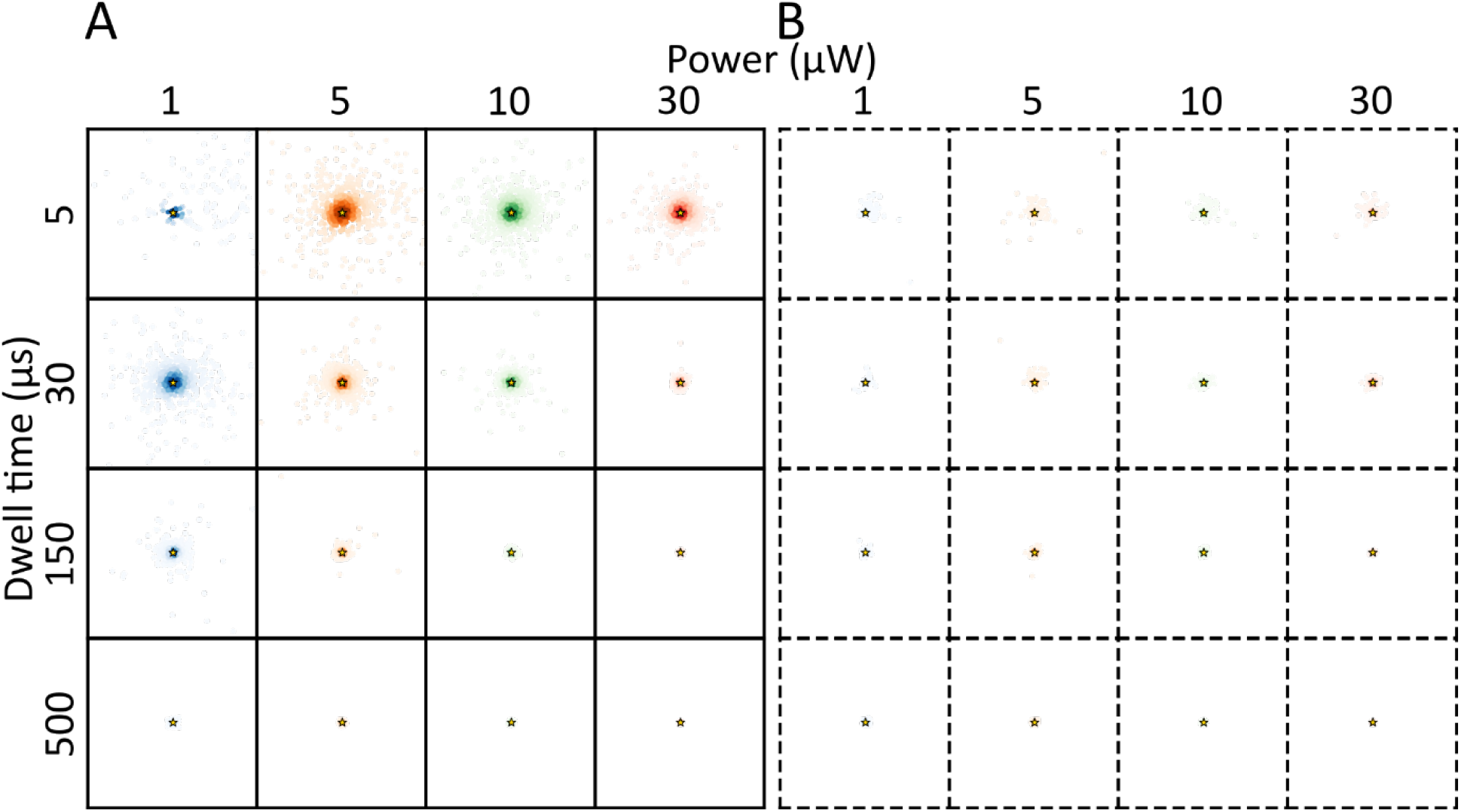
Maps of simulated final estimated locations (coloured dots) for a hypothetical DL755 like fluorophore, with no long-lived redox state, at position (1,1) (golden star) following an iterative MINFLUX localization. The figure shows simulated localization estimates for different powers, different beam dwell times and without (A) and with (B) pattern repeats (dotted boxes, pattern repeat of 5). With pattern repeats, most of the estimates are very close to the actual position with similar localization errors between different powers and beam dwell times. The ROI (each box) has the dimensions -100nm to +100nm in the x and y directions. The difference in color transparency shows the density of estimated locations, calculated using a kernel density estimate(KDE) with a Gaussian kernel.

Notably, alleviating photo-isomerization effects in this way would also synergistically improve photon counts and signal-to-background conditions, as long as higher excitation intensities and doses will not cause photobleaching or transitions into other, more long-lived dark states. In our simulations, we did not include photobleaching, since this was not directly observed within the relaxation times covered in the TRAST measurements (up to 10ms), and since typical emission ON times of single molecules in MINFLUX experiments are much longer (10ms or longer) than the dwell and iteration times simulated. Similar results as in Figure 4 were found also for corresponding simulations of a hypothetical isomerization-only AF647 (SI Section S15, Figure S15), further suggesting that buildup into the 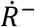 state is a major reason for blinking-induced localization errors of cyanines in MINFLUX experiments. However, it should be noted that 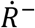 state formation is also intimately coupled to the slow (milliseconds-seconds), low duty cycle switching required for the single-molecule localization, as used in STORM imaging.^13^ Thus, if 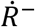 state suppression is used as a strategy to improve localization, then other means will be required to perform the slow time-scale fluorophore switching.

One means to replace the intrinsic (slow) switching of fluorophores is to instead generate the switching by the point accumulation for imaging in nanoscale topography (PAINT) method.^37^ By using PAINT, and transient binding of fluorophore-labeled single-stranded DNA (DNA imager strands) to complementary strands (DNA docker strands) attached to the target molecules (DNA-PAINT),^38^ several constraints on the fluorophores and their switching kinetics can be relieved in MINFLUX experiments.^39^ Cyanine fluorophores, requiring certain imaging buffers to attain appropriate slow switching properties, may then be replaced by other fluorophores. Alternatively, for NIR MINFLUX imaging, and since most available fluorophores in the NIR are cyanines, another strategy is to stay with cyanine fluorophores, use DNA-PAINT, and to modify the imaging buffers to suppress 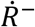 transitions and increase their brightness, without concern about the intrinsic slow switching properties of the fluorophores.

As a first step, to realize this latter strategy, we added ascorbic acid (AA 1mM) and methyl viologen (MV 1mM) as a reducing and oxidizing (ROXS)^34^ system into the fluorophore samples. TRAST curves recorded from DL755 in a deoxygenated TAE (Tris, acetic acid and EDTA) buffer with GODCAT then show an almost complete suppression of 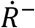 buildup (Figure 5A). For AF647, a similar, but somewhat smaller effect from the ROXS system was observed (SI Section S16 Figure S16). In DL755, with the ROXS system added and photoreduction practically eliminated within relevant time scales for MINFLUX localization, DL755 is very similar to the hypothetical isomerization-only DL755 fluorophore considered in the simulations in Figure 4. It can thus be expected to display similar strongly decreased blinking-attributed localization errors. Moreover, apart from a suppressed fluorophore blinking and 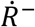 buildup, the added ROXS system has also been found to suppress (irreversible) photobleaching^34^ and protect DNA docker strands from ROS damage.^40^ For typically less photostable NIR fluorophores higher laser irradiances can then be tolerated in MINFLUX experiments, which can further reduce blinking-attributed localization errors (Figure 4), generate higher fluorescence photon numbers, 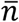, as a basis for the localization, and improve signal-to-background (S-B) conditions. Another, orthogonal strategy to improve fluorophore brightness, particularly in NIR heptamethine cyanines, is to replace water with heavy (deuterated) water in the samples.^26, 41, 42^ Our TRAST experiments showed a two-to three-fold increase in the fluorescence signal of DL755 by use of heavy water (SI Section S17), a corresponding increase in the fluorescence lifetime, and a concomitant decrease in the isomerization relaxation time. These effects are all in agreement with a lower hydrogen bond-mediated quenching of excited singlet states, of both the N and P states.

**Figure 5:**
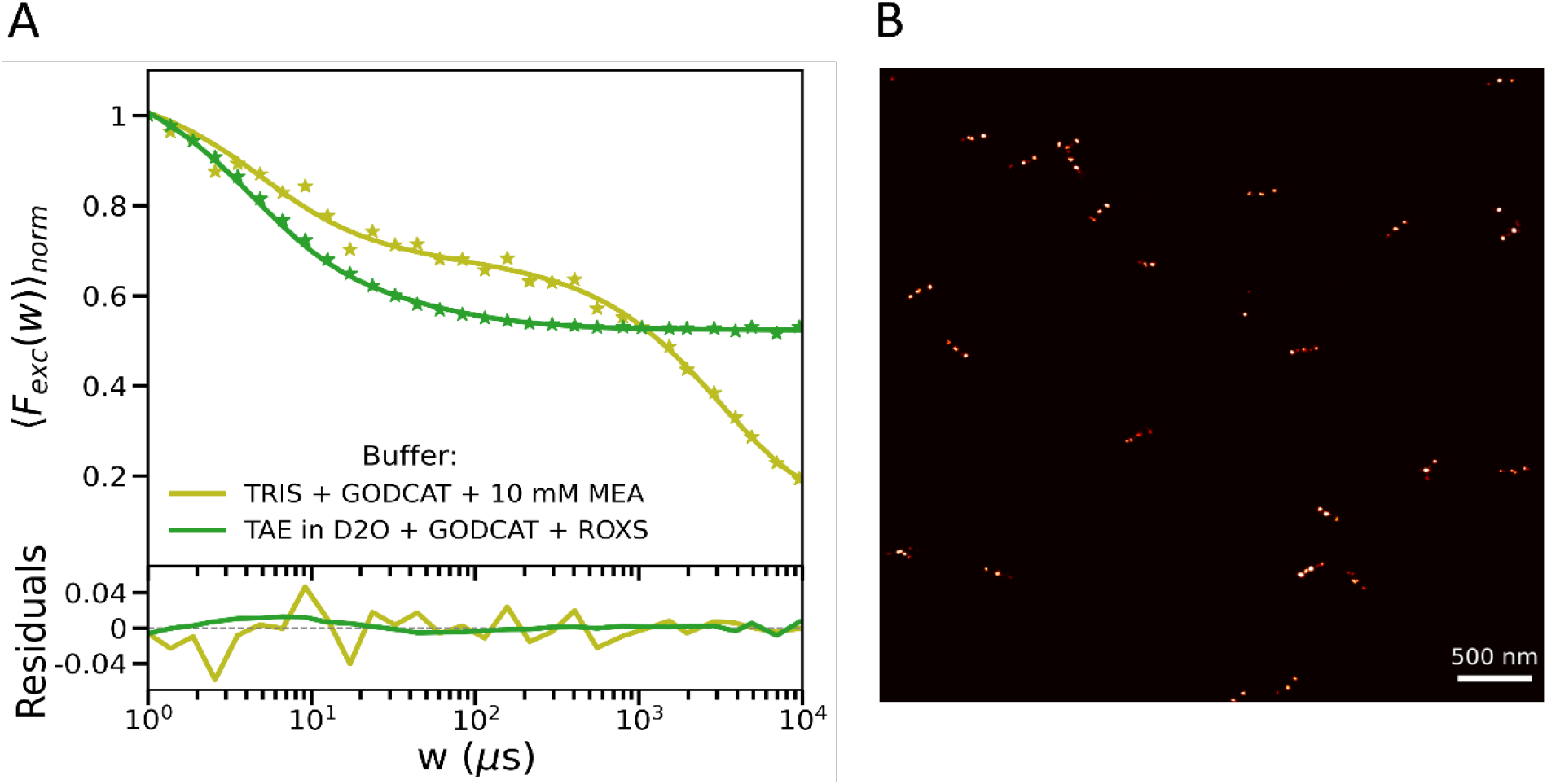
A) TRAST curves recorded for DL755 conjugated to an antibody in a STORM switching buffer (TRIS,GODCAT & MEA) and for DL755 conjugated to imager DNA strand in a deuterated redox buffer (TAE in D_2_0,GODCAT & ROXS)(See Methods for details).B) DNA-PAINT MINFLUX image of DNA origami nanorulers with DNA docker strands located at three positions 80nm apart, and with DL755-labeled complementary imager DNA strands in a deuterated redox buffer (TAE in D_2_0,GODCAT & ROXS), see Methods for details.

### Experimental demonstration of NIR-MINFLUX imaging

Having identified an imaging buffer which suppresses 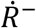 buildup and increases the brightness of DL755 (and other heptamethine cyanines), we next demonstrated this strategy for MINFLUX imaging in practice, at the same time demonstrating MINFLUX extended into the NIR. DNA origami nanorulers (GATTA PAINT 80, GATTAquant) immobilized on a cover glass were imaged by a DNA-PAINT approach, where DNA imager strands with DL755 could bind and then dissociate from complementary DNA strands (docker strands) at three equidistant (80nm) sites on the nanorulers (see Methods section). The sample was kept in the same redox buffer (TAE, GODCAT and ROXS, with deuterated water) as found to suppress 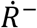 buildup (Figure 5A). The imaging was performed with a MINFLUX platform from Abberior Instruments, modified to incorporate NIR excitation (750nm) and detection (see Methods section). A representative MINFLUX image is shown in Figure 5B. For comparison, we also attempted to image the same sample with a commerically available DNA-PAINT imaging buffer. However, this resulted in images with almost no succesful localizations (data not shown). Additional TRAST measurements confirmed that the 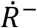 buildup was not suppressed in a commerical imaging buffer, not affecting the redox transitions of DL755 (SI section S18 Figure S18).. Similarly, using AF647 as fluorophore, and combining the same balanced redox buffer as for DL755 (Figure 5A) with DNA-PAINT, also resulted in improved MINFLUX images (SI section 19, Figure S19).

With DNA-PAINT MINFLUX, intrinsic low duty cycle, slow switching properties of the fluorophores are not required, but instead executed via binding and dissociation of DNA-imager strands. Thus, other NIR fluorophores than heptamethine cyanines can also be considered for MINFLUX imaging. Accordingly, we investigated the dark state transitions of the NIR oxazine fluorophore Atto700. For Atto700, no significant photo-induced redox transitions were found in a commercial DNA-PAINT imaging buffer, or in a regular PBS buffer (SI Section S20, Figure S20A). Under these buffer conditions (which differed from those used for the heptamethine cyanines) we could then successfully perform DNA-PAINT MINFLUX imaging, using Atto700-labeled imager strands and the same nanorulers as shown in Figure 5B (SI Section S20, Figure S20B). Similar to the heptamethine cyanines, however, when using a redox buffer in which redox state buildup in Atto700 is prominent (SI section S20, Figure S20A) only very sparse localizations were found in the MINFLUX images (SI section S20, Figure S20C). This further underlines that fluorophore dark state transitions/blinking within typical time ranges for the TCPs is a major factor limiting the localization precisions and probabilities in MINFLUX imaging. On the other hand, it also shows that by suppression of such photophysical transitions, additional fluorophore categories, including NIR fluorophores, become eligible for MINFLUX imaging.

## Conclusions

In this work, we systematically studied the blinking/switching properties of NIR heptamethine cyanine fluorophores and AF647, over a time scale spanning several orders of magnitude (µs-10ms), and under different labeling and sample conditions relevant for MINFLUX imaging. From the acquired data, photodynamic models with rate parameters were determined, and next used in simulations of the fluorophore blinking behavior under representative MINFLUX excitation conditions. This study confirms that faster time scale (µs-ms) blinking of the fluorophores, especially from photo-induced redox state transitions, can lead to significant localization errors and is a major underlying reason for the lowered localization probability experienced in MINFLUX measurements. By use of a balanced redox buffer suppressing such transitions, and replacing the intrinsic slow switching of the fluorophores by a DNA-PAINT approach these problems can be alleviated, allowing NIR-MINFLUX imaging to be realized. Our findings are relevant not only for MINFLUX imaging, but to several other SRM techniques incorporating time-modulated illumination patterns in the localization.^19-21^ They also allow us to define different strategies to minimize these errors, and open for the use of a braoder range of fluorophores in MINFLUX imaging and in SRM in general.

## Methods

### Sample preparation

Stock solutions of Alexa Fluor™ 647 NHS Ester (AF647, Invitrogen, USA), DyLight™ 755 NHS Ester (DL755, Thermo Scientific, USA), Alexa Fluor™ 750 NHS Ester (AF750, Invitrogen, USA), CF™ 750 succinimidyl ester (Sigma-Aldrich, USA) were prepared in DMF and stored at −20 °C and then diluted in different solvents just before measurements to a final concentration of around 500nM.

In the TRAST measurements on antibody-conjugated dyes, all dyes were conjugated to AffiniPure Sheep Anti-Mouse IgG (H+L) (Biozol, Germany) with a degree of labeling (DOL) between 0.1 to 0.2 to be assured that there was just one fluorophore on each antibody.

The imager strand F1 were purchased with dye (Alexa Fluor 647 and ATTO 700) conjugation from Massive Photonics. Dylight 755, IR800CW and Dyomics751 were conjugated to the oligonucleotide in PBS, through a reaction with five-fold excess of NHS conjugated dyes over DNA, with 50mM sodium bicarbonate (NaHCO_3_, pH 9) at RT, and with a shaker (600 rpm) for 2 hours. After reaction, 0.1x volume of 3M Sodium acetate (pH 5.2) and 2.5x volume of ethanol were added and kept at -20 °C for 2 hours, allowing for DNA precipitation. The mixture was then centrifuged at 14,000g for 20 minutes and the supernatant with free dyes was removed. The remaining pellet was then resuspended, washed three times with 70% ethanol, centrifuged at 14,000g for 5 minutes each time, then dried, resuspended and stored at -20 °C in Tris EDTA buffer.

### Buffer solutions

The buffers used in the TRAST and MINFLUX imaging measurements were as follows(all chemicals purchased from Sigma-Aldrich, USA, unless stated otherwise):

#### STORM switching buffer

50mM Tris buffer, 10mM NaCl, GODCAT (10% w/v D-(+)-glucose, 64 µg/ml catalase (C100) and 0.4 mg/mL glucose oxidase (G2133)), 10mM MEA (Cysteamine).

#### ROXS redox buffer

2.5x TAE (Tris, acetic acid and EDTA) in H_2_O or in D_2_O for deutarated buffer, 2M NaCl, GODCAT (1% w/v D-(+)-glucose, 50 µg/ml catalase, 0.6 mg/mL glucose oxidase), ROXS (1mM L-Ascorbic acid, 1mM Methyl viologen dichloride hydrate).

#### Commercial DNA-PAINT imaging buffer

1x Massive Photonics Imaging buffer (Massive Photonics).

#### Trolox redox buffer

2.5x TAE (Tris, acetic acid and EDTA) in H_2_O, 2M NaCl, GODCAT (1% w/v D-(+)-glucose, 50 µg/ml catalase, 0.6 mg/mL glucose oxidase), 5mM Trolox (200mM stocks of (±)-6-Hydroxy-2,5,7,8-tetramethylchromane-2-carboxylic acid was prepared in DMSO and kept in -20ºC for less than 6 months).

### Transient state (TRAST) spectroscopy

TRAST measurements were carried out on a home-built TRAST setup based on an inverted epi-fluorescence microscope. In short, fluorescence is excited by the beam of a diode laser at 638 nm (140 mW, Cobolt, 06-MLD) or at 750 nm (500 mW, MBP communication Inc., Canada), passing appropriate excitation filters (FF01-637/7 (Semrock) and ET740/40x (Chroma) respectively). The 638 nm laser beam was modulated directly by external triggering, and the 750 nm laser beam by an acousto-optic modulator (MCQ110-A1,5-IR, AA Opto Electronics, France). The expanded laser beam was defocused by a convex lens, reflected by a dichroic mirror (ZT405/473/559/635/748rpc-UF3 from Chroma), and then focused close to the back aperture of the objective (alpha Plan-Fluar 100x/1.45 Oil, ZEIZZ, USA) to produce a wide-field illumination in the sample (beam waist ω_0_ = 15–25 μm (1/e^2^ radius)). The fluorescence signal was collected by the same objective, passed through the same dichroic mirror and a filter (638 notch filter and FF02-809/81 (Semrock) emission filter, respectively) to remove scattered laser light, and then fed to a digital sCMOS camera (ORCA-Fusion BT, Hamamatsu Photonics, Japan). The experiments were controlled and synchronized by custom software implemented in Matlab. A digital I/O card (PCI-6602, National Instruments) was used to trigger the camera and generate pulse trains sent to the AOM driver unit.

### Nanoruler sample preparation

DNA origami - based nanorulers (GATTA-PAINT 80, GATTAquant GmbH) with three DNA-PAINT F1 docking sites (Massive Photonics) with 80nm distance between the sites were immobalized in a 8-well glass bottom chamber (ibidi µ slide chamber) according to manufacturer instructions. Briefly, the chamber was washed with PBS three times (3 minutes each) before adding 1 mg/mL BSA-biotin solution in PBS for 5 minutes. After incubation with BSA-biotin, the solution is removed, 3x washed with PBS and incubated with 1mg/mL neutravidin solution for 5 minutes. The chamber is then washed three times with PBS supplemened with 10mM magnesium chloride (so-called immobilization buffer IB). The nanoruler solution is then diluted 1:3 with IB and incubated in the chamber for 20 minutes.The chamber is again washed three times with IB. 100uL of undiluted nanorod dispersion (A12-40-980-CTAB-DIH-1-25, Nanopartz Inc.) were then incubated for 20 minutes for stabilization of samples during MINFLUX imaging. The chamber were then washed three times with IB and replaced with appropriate imaging buffers with imager strands and with 20mM MgCl_2_ for stabilization of DNA nanorulers. DNA-PAINT MINFLUX was performed with µ slides in air saturated conditions, since the photophysical environment provided by the redox buffer was measured to be preserved for at least for 3 hours (SI section S21).

### MINFLUX imaging

MINFLUX imaging was performed with a platform from Abberior Instruments (see ^36^), prototyped with a NIR excitation laser (750nm, 500 mW, MBP communication Inc., Canada) and with a single photon avalanche detector (SPAD) with enhanced NIR detection (Excelitas, SPCM-NIR). Data acquisition and instrument control was done using the Imspector software (v. 16.3.16297M-minflux, Abberior Instruments) and Imspector sequence Imaging2D (SI section S22) was used for 2D imaging. A region-of-interest (ROI) in the samples was selected, displaying sufficient blinking upon confocal imaging. Once a ROI was selected, the stabilization system was activated and the starting excitation power (power for the first iteration) was selected before starting the MINFLUX measurements. For the ATTO700 measurements, laser excitaiton at 640nm was used, with a starting laser power of ∼210 µW (measured at the back aperture of the objective)., On the detection side, a regular SPAD (Excelitas, SPCM-AQRH-13-FC) was used with emission filter (685-730nm) and a pinhole size of 0.83AU. For the NIR heptamethine cyanine measurements, a 750nm laser was used, with a starting laser power of ∼180 µW, a NIR-SPAD (Excelitas, SPCM-780-14-FC) for detection (emission filter 770-855 nm) and a pinhole size of 1.1AU. The MINFLUX localizations, obtained from the Imspector software were drift corrected using COMET (Cost-function optimized maximal overlap drift estimation) and rendered in python with a bin size of 5 nm.

### Fluorescence lifetime measurements

Fluorescence lifetime measurements were performed by time-correlated single photon counting (TCSPC), see SI Section S3 for details.

### Simulations of MINFLUX localizations of individual fluorophores

Simulations are described in main text, and with further details in SI, Section S9.

