## Supplementary Information for "Near-infrared MINFLUX imaging enabled by suppression of fluorophore blinking"

Chinmaya Venugopal Srambickal^1,*^, Hanie Esmaeeli^1,*^, Joachim Piguet^1^, Lenny Reinkensmeier^2^, René Siegmund^2^, Mark Bates^2^, Alexander Egner^2^, Jerker Widengren^1,**^

^1^ Experimental Biomolecular Physics, Bio-Opto-Nano Unit, Department of Applied Physics, Royal Institute of Technology, SE-10691 Stockholm, Sweden

^2^ Department of Optical Nanoscopy, Institute for Nanophotonics, D-37077 Göttingen, Germany

^*^ Contributed equally

**Section S1:** TRAST concept

In TRAST spectroscopy/imaging, the population dynamics of photoinduced, non- or weakly fluorescent, long-lived transient states of fluorescent molecules, such as triplet, photo-isomerized and photo-redox states, are determined from the average fluorescence intensity detected in the sample, when subject to different excitation pulse trains. ^1, 2^

Considering a homogeneous fluorophore sample, with concentration, *c*, subject to a rectangular excitation pulse with a constant excitation photon flux, $\Phi_{\mathrm{exc}}$, starting at $t=0$, the recorded fluorescence intensity can be described by

$F\left( t \right)=c\cdot{{}^{1}q}_{F}\cdot{{}^{1}q}_{D}{\cdot\sigma}_{1}\iiint\left( CEF\left( \bar{r} \right)\cdot\Phi_{\mathrm{exc}}\left( \bar{r} \right)\cdot\sum_{j=1}^{n} \left( Q_{j}\left[ A_{j} \right]\left( \bar{r},t \right) \right) \right)dV$	(S1)

Here, $\left[ A_{j} \right]$ denotes the population probability of the i:th photophysical state, ${{}^{1}q}_{F}$ and ${{}^{1}q}_{D}$ denote the fluorescence quantum yield of state A_1_ and the overall detection quantum yield of its emission, respectively. $\sigma_{1}$ is its excitation cross section and $CEF\left( \bar{r} \right)$ is the collection efficiency function of the setup. With A_1_ representing the brightest photophysical state, $Q_{j}$ then denotes the relative brightness of the other states (*j*=2,…n), compared to that of A_1_, and with $Q_{1}$ (the brightness of A_1_) normalized to one.

At onset of excitation, equilibration between any ground and excited singlet states typically occur with a relaxation time in the nanosecond time range. This so-called anti-bunching relaxation time is typically much faster than the characteristic relaxation(s) of the more long-lived dark transient states. The anti-bunching can thus be disregarded on a $\mu s$ to ms time scale after onset of excitation, at which relaxations into dark or weakly emissive transient state often take place. Similar relaxations can also be observed in the time-averaged fluorescence signal resulting from a rectangular excitation pulse of duration $w$

$\left\langle F_{\mathrm{exc}}\left( w \right) \right\rangle=\frac{1}{w}\int_{0}^{w} F\left( t \right) dt$	(S2)

when $w$ is increased from the $\mu s$ to the ms time range. Analyzing how $\left\langle F_{\mathrm{exc}}\left( w \right) \right\rangle$ varies with $w$ then allows the population kinetics of long-lived photo-induced states of the fluorophore to be determined, which is the general basis for TRAST monitoring.

To obtain sufficient photon counts, even for short $w$, the fluorescence intensity resulting from an excitation pulse train of *M* identical pulse repetitions is typically recorded. *M* is adjusted to maintain a constant laser illumination time, $t_{ill}=M\cdot w$, for all $w$. A so-called TRAST curve is then produced by calculating the time-averaged fluorescence signal during excitation for each pulse train, normalized for a given pulse duration, $w_{0}$

$\left\langle F_{\mathrm{exc}}\left( w \right) \right\rangle_{\mathrm{norm}}=\left( \frac{1}{M}\sum_{i=1}^{M} \left\langle F_{\mathrm{exc}}\left( w \right) \right\rangle_{i} \right)/\left( \frac{1}{M_{0}}\sum_{i=1}^{M_{0}} \left\langle F_{\mathrm{exc}}\left( w_{0} \right) \right\rangle_{i} \right)$	(S3)

The pulse duration used for normalization, $w_{0}$, is chosen to be short enough (typically sub-μs) not to lead to any noticeable build-up of dark transient states, yet longer than the nanosecond anti-bunching rise time of $F\left( t \right)$ upon onset of excitation. *M*_0_ refers to the number of pulses used in the normalization.

In the above expression, $\left\langle F_{\mathrm{exc}}\left( w \right) \right\rangle_{i}$ represents the total signal collected from the *i*:th pulse in a pulse train, as defined in Eq. S2. By using a low excitation duty cycle, in this work $\eta=0.001-0.01$, fluorophores are allowed to fully recover back to the all-*trans* ground singlet state, S_0,_ before the onset of the next pulse. In the normalization step of Eq. S3, several parameters used to calculate $F\left( t \right)$ in Eq. S1 cancel out. The final expression for $\left\langle F_{\mathrm{exc}}\left( w \right) \right\rangle_{\mathrm{norm}}$ therefore becomes independent of $c$ as well as of the absolute $q_{D}$ and $q_{F}$ values for the emissive species:

$\left\langle F_{\mathrm{exc}}\left( w \right) \right\rangle_{\mathrm{norm}}=\frac{\int_{t=0}^{w} \left( \iiint\left( CEF\left( \bar{r} \right)\cdot\Phi_{\mathrm{exc}}\left( \bar{r} \right)\cdot\sum_{i=1}^{n} \left( Q_{i}\left[ A_{i} \right]\left( \bar{r},t \right) \right) \right)dV \right)dt}{w\iiint\left( CEF\left( \bar{r} \right)\cdot\Phi_{\mathrm{exc}}\left( \bar{r} \right)\cdot\sum_{i=1}^{n} \left( Q_{i}\left[ A_{i} \right]\left( \bar{r},0 \right) \right) \right)dV}$	(S4)

**Section S2:** TRAST curves of AF750 and CF750 recorded at different excitation intensities

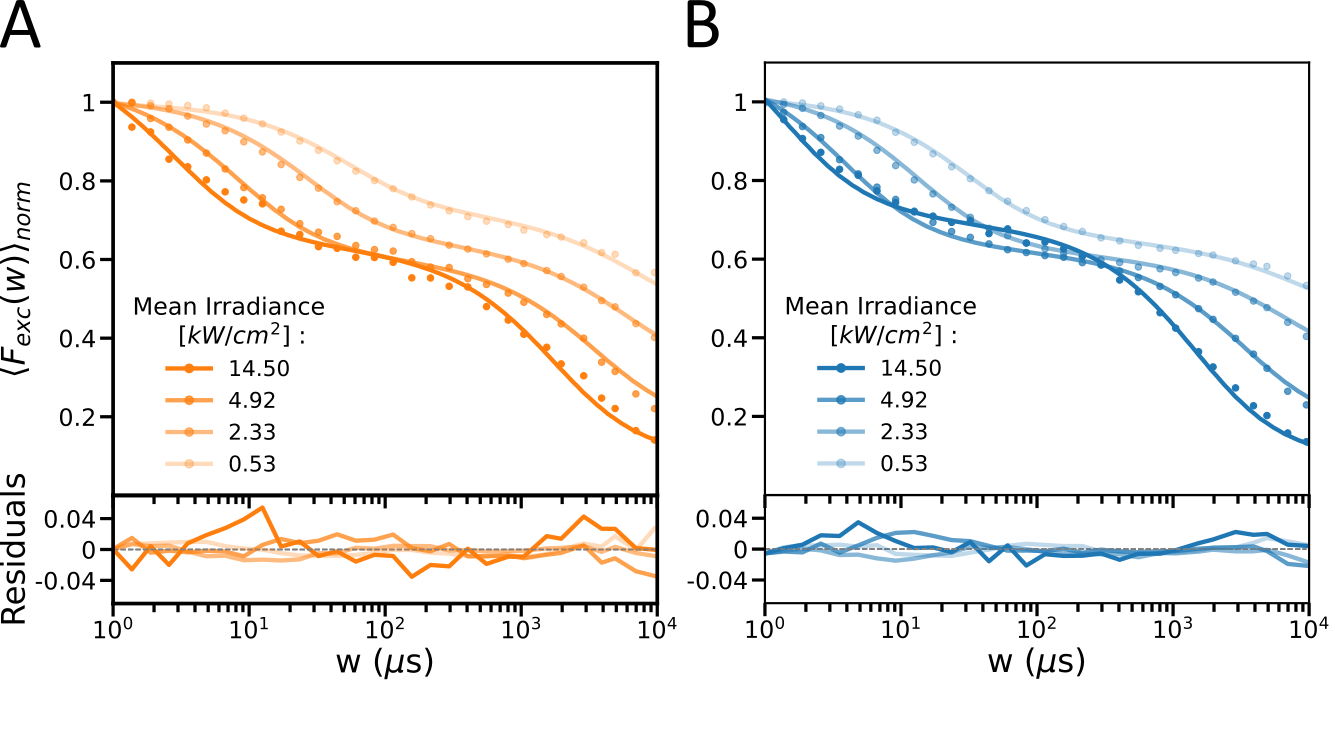

**Figure S1:** TRAST curves measured from free A) CF750 and B) AF750 in PBS, with different (750nm) excitation intensities applied. Dots: experimental data, lines: fitted TRAST curves (see main text), fitting residuals below.

**Section S3:** Fluorescence lifetime measurements

Fluorescence lifetime measurements were performed by time-correlated single photon counting (TCSPC) using a commercial, epi-illuminated, confocal laser scanning microscope (Olympus FV1200). Fluorophore solution samples were excited by the focused beam of a 640 nm diode laser (LDH-D-C-640 , PicoQuant GmbH, Berlin) and of a 750 nm RF tunable laser (SuperK SELECT from NKT-photonics) operated in pulsed mode. The emitted fluorescence was collected back through the microscope objective (UPlanSApo 60x/1.2W, Olympus), passed through a dichroic mirror (ZT405/488/635rpc-UF2, Chroma or T770lpxr-UF2, Chroma), an emission filter (HQ720/150, Chroma, or 809/81 Brightline, Semrock, Semrock), and focused onto a pinhole (50µm diameter) in the back focal plane. The fluorescence signal was finally split and directed on two avalanche photodiodes (Tau-SPAD, PicoQuant GmbH, Berlin). Instrument response functions (IRFs) were determined from the back-reflected light from the laser excitation pulses. The signals were fed into a data acquisition card (Hydraharp 400, Picoquant GmbH), deconvoluted and then fit to an exponential decay based on non-linear least squares minimization (Symphotime, Picoquant GmbH).

**Section S4:** Photophysical model and rate equations for the cyanine fluorophores

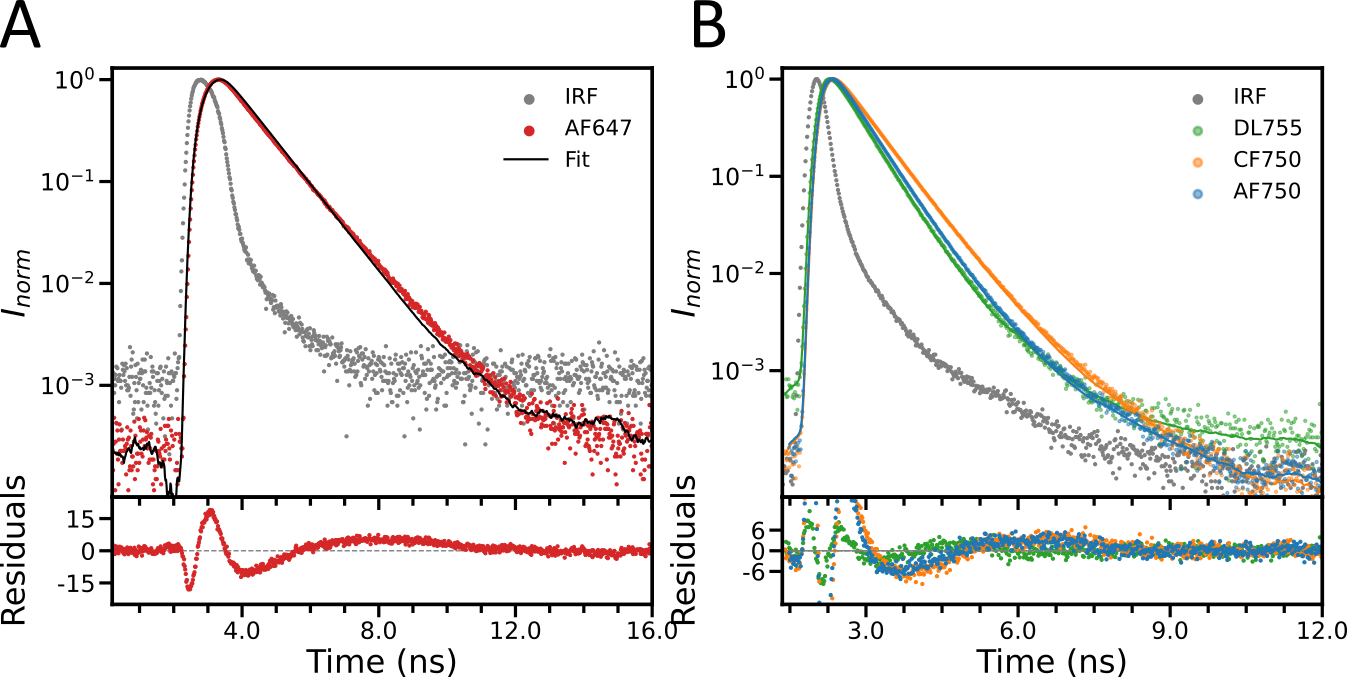

**Figure S2**: Time-correlated single photon counting (TCSPC) measurements data from A) free AF647 with 640 nm excitation and B) NIR dyes with 750nm excitation in PBS. The fitted curve for AF647 shows a lifetime of 1.07 ns, whereas the lifetime fitted for the NIR fluorophores, DL755, CF750 and AF750, are 0.47 ns, 0.6 ns and 0.51 ns, respectively.

With the photophysical model shown in Figure 1D (main text), the population probabilities of the different states of the cyanine fluorophores, subject to a constant excitation photon flux of $\Phi_{exc}$ starting at time t=0, is given by

$\frac{d}{dt}\bar{A}\left( t \right)=M\cdot\bar{A}\left( t \right)$	(S5)

Here, $\bar{A}\left( t \right)=\left[ \left[ N \right]\left( t \right), \left[ P \right]\left( t \right), \left[ T \right]\left( t \right), [\dot{R}^{-}](t) \right]^{T}$ represents the population probabilities of the all-*trans*, the photo-isomerized, triplet and photo-reduced states of the fluorophore and

$M=\left[ \begin{matrix} \begin{matrix} {-(k}_{iso}´+k_{isc}´) & k_{biso}´ \\ k_{iso}´ & {-k}_{biso}´ \end{matrix} & \begin{matrix} k_{T} & k_{ox} \\ 0 & 0 \end{matrix} \\ \begin{matrix} k_{isc}´ & 0 \\ 0 & 0 \end{matrix} & \begin{matrix} -(k_{T}+k_{red}) & 0 \\ k_{red} & -k_{ox} \end{matrix} \end{matrix} \right]$	(S6)

is the rate matrix describing the transitions between the states. In the model, it can be assumed that equilibration between the ground and excited singlet states of N and P take place on a much faster time scale than the relaxation of $\bar{A}\left( t \right)$, i.e. the time scale of the fluorophore dark state transitions and the TRAST experiments (1µs-10ms), and at which also the MINFLUX beam localization procedure typically operates. [*N*] and [*P*] thus denote the total probabilities of the fluorophore to be in either its ground or excited singlet state for N and P, respectively. In the matrix, we can then also assign effective isomerization ($k_{iso}´$), back-isomerization ($k_{biso}´$) and intersystem crossing ($k_{isc}´$) rates. The effective isomerization rate, from N to P is given by:

$k_{iso}´=k_{iso}\cdot\frac{\sigma_{N}\cdot\Phi_{exc}}{\sigma_{N}\cdot\Phi_{exc}+k_{10}^{N}}$	(S7)

with $\sigma_{N}$ denoting the excitation cross sections of the singlet ground state of N, and $k_{10}^{N}$ signifying the decay rate from the excited singlet state to the ground singlet state in N. Since $k_{10}^{P}$ and $\sigma_{P}$ could not be individually determined, we defined the back-isomerization rate from P to N by:

$k_{biso}´=k_{biso}\cdot\frac{\sigma_{P}\cdot\Phi_{exc}}{\sigma_{P}\cdot\Phi_{exc}+k_{10}^{P}}+k_{biso}^{Th}=\left\{ k_{10}^{P}\gg\sigma_{P}\cdot\Phi_{exc} \right\}=\sigma_{biso}\cdot\Phi_{exc}+k_{biso}^{Th}$	(S8A)

, where $k_{biso}^{Th}$ denotes the thermal back-isomerization rate, and where the back-isomerization cross section is defined as:

$\sigma_{biso}=k_{biso}\cdot\frac{\sigma_{P}}{k_{10}^{P}}$	(S8B)

Analogous to Eq. S3, the effective intersystem crossing rate from N to T can be defined as:

$k_{isc}´=k_{isc}\cdot\frac{\sigma_{N}\cdot\Phi_{exc}}{\sigma_{N}\cdot\Phi_{exc}+k_{10}^{N}}$	(S9)

The initial condition for Eq. (S1) is

$\bar{A}\left( 0 \right)=\left[ 1 0 0 0 \right]^{T}$	(S10)

, based on the finding that, in absence of excitation, polymethine cyanine fluorophores typically exist in their all-*trans* state.^3^ All fluorophores can thus be assumed to be in the all-*trans* singlet (ground) state before onset of excitation at $t=0$.

For a rectangular excitation pulse, $\Phi_{exc}$ is constant throughout the excitation duration and the matrix $M$ is not time dependent. The general solution to Eq S1 is then

$\bar{A}\left( t \right)=e^{Mt}\cdot\bar{A}\left( 0 \right)$	(S11)

The dependence of the detected fluorescence at time, *t*, after onset of excitation is then given by

$F\left( t \right)={}^{1}{q_{F}\cdot{}^{1}{q_{D}}\cdot k_{10}^{N}}\cdot\frac{\sigma_{N}\cdot\Phi_{exc}}{\sigma_{N}\cdot\Phi_{exc}+k_{10}^{N}}\cdot\left[ N \right]\left( t \right)+{}^{2}{q_{F}\cdot{}^{2}{q_{D}}\cdot k_{10}^{P}\cdot}\frac{\sigma_{P}\cdot\Phi_{exc}}{\sigma_{P}\cdot\Phi_{exc}+k_{10}^{P}}\cdot\left[ P \right]\left( t \right)$	(S12)

, with $\sigma_{N}$ and $\sigma_{P}$ denoting the excitation cross sections of the N and P state, respectively. ${}^{X}{q_{F}}$ is the fluorescence quantum yield and ${}^{X}{q_{D}}$ the overall detection quantum yield of the emission from N (X=1) and P (X=2) state, respectively. For the excitation conditions in our study, $k_{10}\gg\sigma_{N}\cdot\Phi_{exc}, \sigma_{P}\cdot\Phi_{exc}$, so that we can assume

$F\left( t \right)={}^{1}{q_{F}\cdot{}^{1}{q_{D}}\cdot\sigma_{N}\cdot\Phi_{exc}\cdot}\left( \left[ N \right]\left( t \right)+Q\cdot\left[ P_{2} \right]\left( t \right) \right)$	(S13)

, with $Q=({}^{2}{q_{F}\cdot{}^{2}{q_{D}}\cdot\sigma_{N})/({}^{1}{q_{F}\cdot{}^{1}{q_{D}}\cdot\sigma_{P_{2}})}}$ representing the relative brightness of P, compared to N.

**Section S5:** Calculated TRAST curves as a result of the photophysical model

With the transient state population kinetics of the cyanine fluorophores described by the four-state photophysical model shown in Figure 1b, $\bar{A}\left( t \right)=\left[ \left[ N \right]\left( t \right), \left[ P \right]\left( t \right), \left[ T \right]\left( t \right), [\dot{R}^{-}](t) \right]^{T}$ can be assigned to represent the population probabilities of the all-*trans*, the photo-isomerized, triplet and photo-reduced states of the fluorophore, respectively. For the studied cyanine fluorophores, N and the P represent emissive states, where $\left[ N \right]$ and $\left[ P \right]$ represent the total population probability of N and P, to be in either their ground or excited singlet states. For a fluorophore subject to a rectangular excitation pulse with constant $\Phi_{\mathrm{exc}}$ starting at $t=0$, the fluorescence intensity response then reflects the time dependence of $\left[ N \right]$ and $\left[ P \right]$:

$F\left( t \right)=c\cdot{{}^{1}q}_{F}\cdot{{}^{1}q}_{D}{\cdot\sigma}_{N}\iiint\left( CEF\left( \bar{r} \right)\cdot\Phi_{\mathrm{exc}}\left( \bar{r} \right)\cdot\left( \left[ N \right]\left( \bar{r},t \right)+Q\cdot\left[ P \right]\left( \bar{r},t \right) \right) \right)dV$	(S14)

Here, Q is the relative brightness of P compared to N. ${{}^{1}q}_{F}$ and ${{}^{1}q}_{D}$ denote the fluorescence quantum yield and the overall detection quantum yield of the emission from the excited singlet state of N. $\sigma_{N}$ denotes the excitation cross section of the ground singlet state of N.

With all fluorophores in their all-*trans* (N) state at onset of excitation ($\bar{A}\left( 0 \right)=\left[ 1 0 0 0 \right]^{T}$, see Eq. S10), the averaged and normalized fluorescence intensities in the recorded TRAST curves can then be written:

$\left\langle F_{\mathrm{exc}}\left( w \right) \right\rangle_{\mathrm{norm}}=\frac{\int_{t=0}^{w} \left( \iiint\left( CEF\left( \bar{r} \right)\cdot\Phi_{\mathrm{exc}}\left( \bar{r} \right)\cdot\left( \left[ N \right]\left( \bar{r},t \right)+Q\cdot\left[ P \right]\left( \bar{r},t \right) \right) \right)dV \right)dt}{w\iiint\left( CEF\left( \bar{r} \right)\cdot\Phi_{\mathrm{exc}}\left( \bar{r} \right) \right)dV}$	(S15)

The time dependence of $\left[ N \right]$ and $\left[ P \right]$ for a fluorophore subject to a rectangular excitation pulse with constant $\Phi_{\mathrm{exc}}$ starting at $t=0$ is described by Eqs. S5-S13 in section S2.

If $\left[ N \right]$ and $\left[ P \right]$ can be assumed to be constant within the detection volume at any specific time, *t*, during a rectangular excitation pulse, then Eq. S15 simplifies to:

$\left\langle F_{\mathrm{exc}}\left( w \right) \right\rangle_{\mathrm{norm}}=\frac{1}{w}\int_{t=0}^{w} \left( \left[ N \right]\left( t \right)+Q\cdot\left[ P \right]\left( t \right) \right)dt$	(S16)

**Section S6:** TRAST data analysis

The data analysis was performed similarly to in previous work,^4-8^ adapted to the experimental conditions, samples and models used here (see also Eqs. S5-S16).

A complete TRAST experiment consisted of a stack of 30 fluorescence images. Each image represents the total fluorescence signal from an entire excitation pulse train, captured using a camera exposure time of $t_{\exp}={t_{ill}}/\eta$. Images were recorded applying different pulse durations, $w$, distributed logarithmically between 1 µs and 10 ms. They were measured in a randomized order to avoid bias due to time effects. An additional 10 reference frames, all using 1 µs pulse duration to avoid dark state build-up, were inserted at regular intervals between the 30 main images to track any permanent fluorescence photobleaching of the sample.

The TRAST data were analyzed using a software implemented in Matlab, as previously described.^4-8^ The recorded TRAST data was first preprocessed by subtraction of the static ambient background and corrected for photobleaching, as described above.

Averaged fluorescence signals, as used in the generation of the TRAST curves (Eq. S3), were calculated within a region of interest (ROI) corresponding to a ~5 *μ*m radius in the focusing plane on the sample, centered on the excitation beam. Since the excitation beam and the excitation photon flux, $\Phi_{\mathrm{exc}}(\bar{r})$, is not fully uniform, a spatial dependence can be expected on the excitation rates and the resulting electronic state populations. The total fluorescence signal on each pixel of the camera then becomes a convolution of $\left[ N \right]\left( \bar{r},t \right)+Q\cdot\left[ P \right]\left( \bar{r},t \right)$ and the microscope collection efficiency function, $CEF(\bar{r})$. By simulating the whole 3D sample volume, and computing the projected 2D image on the camera, it has been found that pre-computing an average observed excitation rate, $\hat{k}_{01}$, for each ROI to be analyzed, speeds up the fitting significantly, without appreciable loss of accuracy.^4, 7, 8^ An approximate $\hat{k}_{01}$ could thus be computed once, before fitting starts, by weighting $k_{01}(\bar{r})$ by brightness and collection efficiency, $CEF(\bar{r})$:

$\hat{k}_{01}=\frac{\iiint k_{01}\left( \bar{r} \right)\cdot\hat{S}_{1}\left( \bar{r} \right)\cdot CEF\left( \bar{r} \right)dV}{\iiint\hat{S}_{1}\left( \bar{r} \right)\cdot CEF\left( \bar{r} \right)dV}$ (S17)

Here, $\hat{S}_{1}\left( \bar{r} \right)=k_{01}(\bar{r})/(k_{10}+k_{01}(\bar{r}))$ represents the population of excited singlet state fluorophores when in an all-*trans* form, N, at onset of excitation, after equilibration between the ground and excited singlet states of N, but before build-up of the other states.

Fitting of photophysical rate parameters was then performed by simulating theoretical TRAST curves using Eqs. S1−S9 and comparing them to the experimental data. The set of rate parameter values best reproducing the experimental data was then found using nonlinear least-squares optimization. In the fit, the excited-state lifetime, $\tau_{f}$, of N, was fixed to its fitted value determined by the TCSPC measurements and with $1/{\tau_{f}}$ comprising all deactivation rates from the excited state of N.

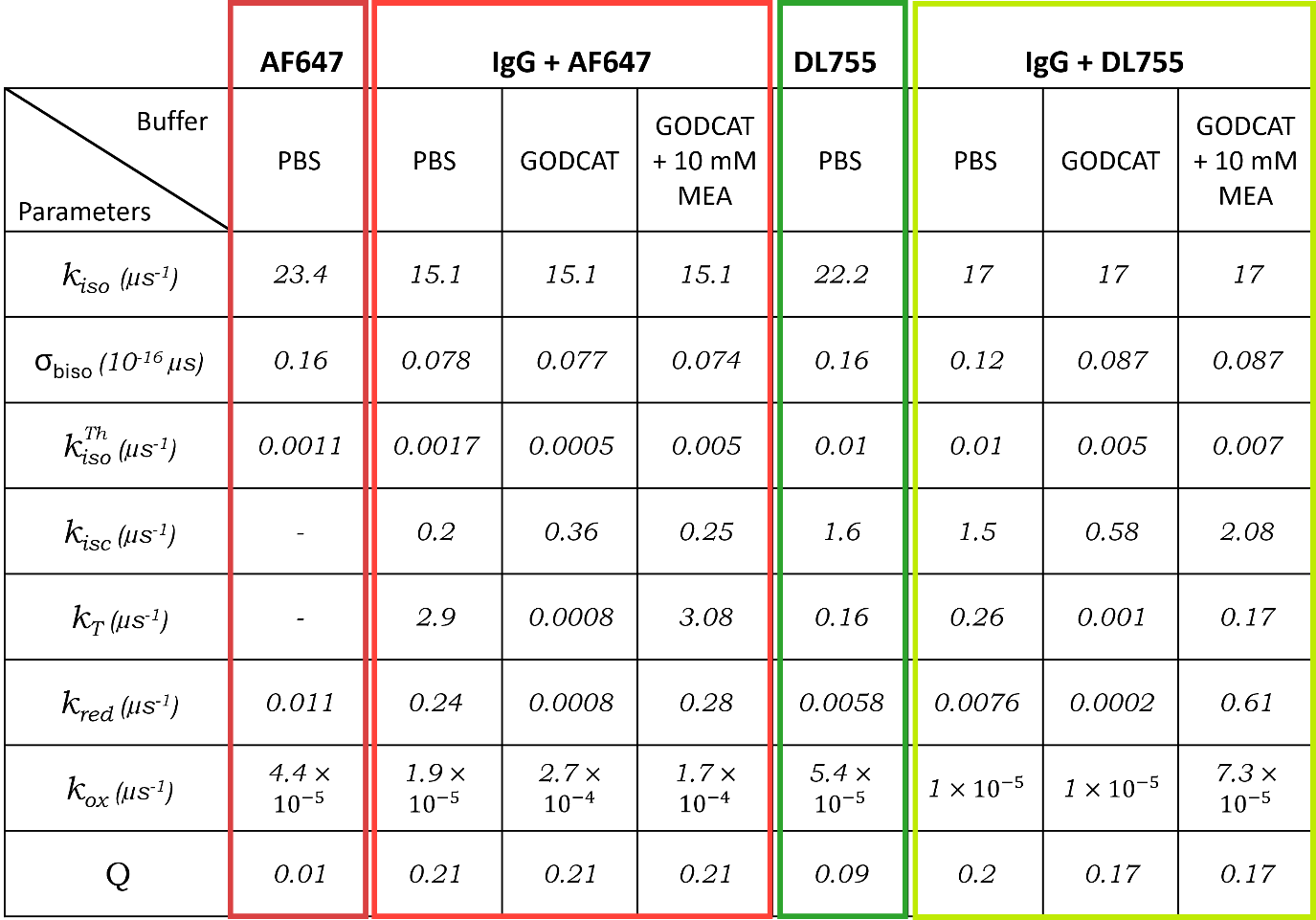
**Section S7:** Fitted rate parameters from TRAST experiments

Table S1: Fitted parameter values from the TRAST experiments. Q values refer to the relative brightness of the P versus the N state, as measured with the multiple notch filter for the 638 excitation and with a 770-850 nm filter for DL755 (750nm excitation).

**Section S8:** TRAST curves recorded from A750 and CF750, conjugated to antibodies, and in presence of MEA and GODCAT

**Figure S3:** TRAST curves recorded from A) CF750 and B) AF750, in free form or when conjugated to an antibody (IgG).TRAST curves recorded from C) CF750 and D) AF750 in a PBS buffer, in a Tris buffer upon deoxygenation (by adding GODCAT), and upon adding MEA. All measurements were performed with excitation at 750nm, 4.7 kW/cm^2^.

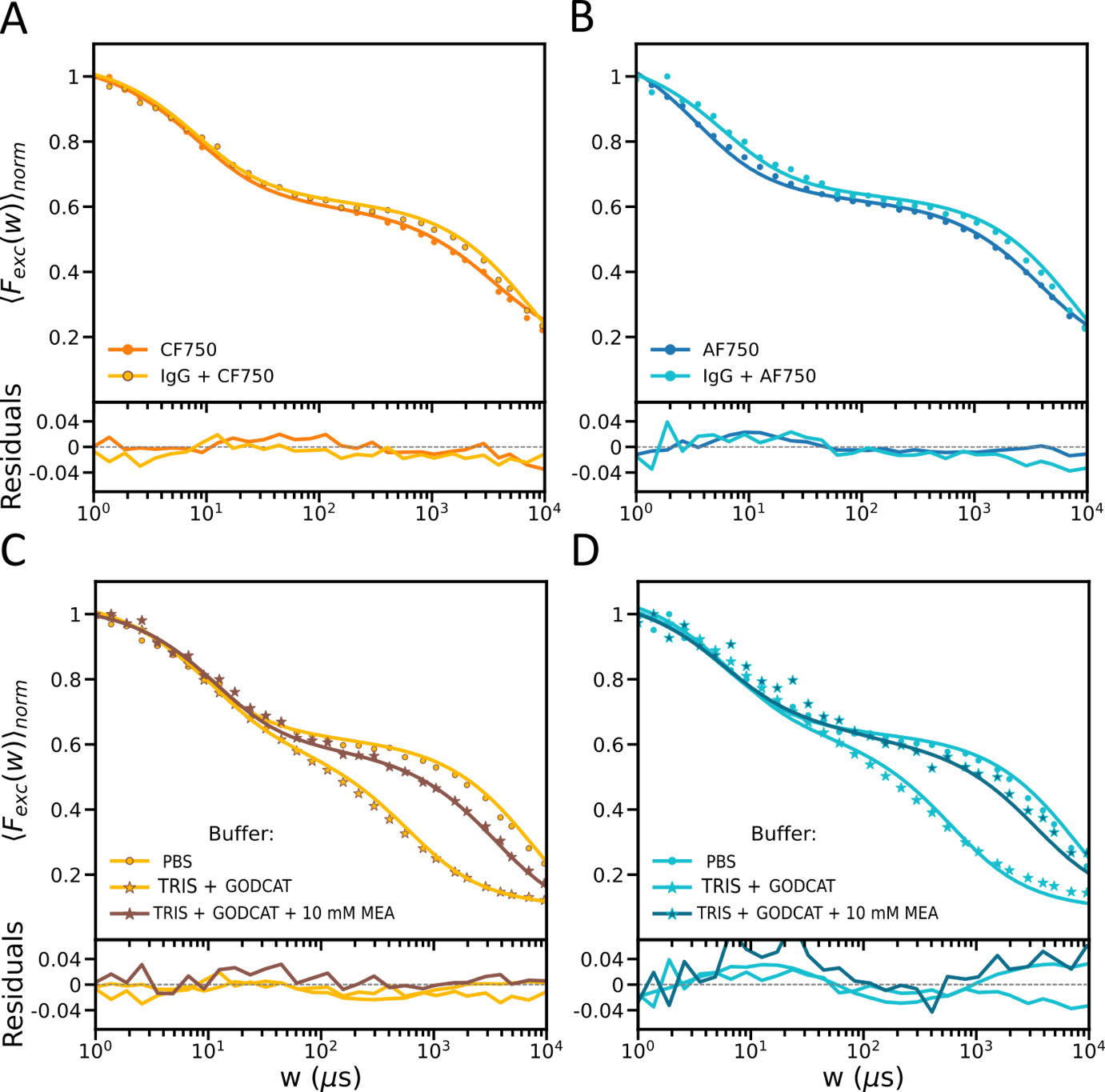

**Section S9:** Simulations of MINFLUX localizations

A program was written in Python to simulate effects of fluorophore blinking behaviour on MINFLUX localization. The simulations consider iterative MINFLUX localizations on a fluorophore at a given location, $\bar{r}_{m}$, while the fluorophore photophysical state evolution is tracked simultaneously. Firstly, the beam positions within each MINFLUX iteration are defined with a targeted coordinate pattern (TCP) consisting of 6 positions in a hexagonal grid with an additional starting position at the center (with the total number of beam positions within each TCP, $K=7$). The diameter ($L$) of the TCP is varied to be 288 nm, 150m, 75nm and 40 nm for iteration 1,2,3, and 4, respectively. To keep the simulation as general as possible, without missing major features of the localization procedure which may be affected by the fluorophore blinking, the pre-localization step (iteration 0) is not simulated, given that there are several different ways of pre-localizing the fluorophore towards the center of the TCP.^9, 10^ In the simulations, we thus assumed that the fluorophore is pre-localized to be within 50nm from its actual position and that the photophysical evolution of the fluorophore due to the pre-localization step is negligible. A region of interest (ROI) of 300 nm x 300 nm was simulated with 1nm sampling, with the TCP center located at (0,0) at the start of simulation. Thus, we selected fluorophore positions within (-50, 0), (0, 50), (0, -50) and (0, 50) and considered at the onset of iteration 1, the fluorophore to be in its N (emissive *trans* isomer) state, according to the initial condition at *t*=0 (Eq. S10). In the simulations we further assumed that only the N state (and not the photo-isomerized P state) is fluorescent.

During the iteration, the beam is placed at each of the beam positions for a beam dwell time, $t_{dwell}$. In case of pattern repeat, the pattern is repeated (within the iteration) such that $t_{dwell}$ is the cumulative time spend at each beam position over all repeats within the TCP iteration. For example, for $t_{dwell}$ = 150µs, pattern repeat = 1, the beam is placed at beam position *i* for 150 µs before it is moved to position *i+1*, whereas for $t_{dwell}$ = 150µs, pattern repeat = 5, the beam is placed for 30µs at position *i* before moving to *i+1*. Once the beam has been placed on each of the $K$ beam positions, the pattern is repeated 4 more times such that the total iteration time is the same in both cases, i.e., $t_{dwell}\times K$.

As the beam is moved from one TCP position to the next within an iteration, the intensity (and the excitation photon flux,$\Phi_{exc})$ experienced by the fluorophore, and hence the excitation rates experienced by the fluorophore at position ($\bar{r}_{m})$ change. This will influence the state evolution, as described in Section S4. Here, $\Phi_{exc}$ is given by the power of the laser beam, its cross section, and how the laser beam is located with respect to the fluorophore. In the simulations, the donut size related parameter (FWHM) was set to 360 nm such that the peak-to-peak diameter is at 1.2FWHM, i.e., 432 nm (See supplementary equation S17 in reference ^11^)). The time and intensity-dependent state evolution of the fluorophore are simulated with time steps ($\Delta t$) of 100 ns. The selected$\Delta t$ is long enough that equilibration between singlet ground and excited states in N and P has taken place (see main text), yet short enough to resolve the µs-ms state evolution. The photophysical state evolution was modelled as a Markovian chain^12, 13^ based on the model obtained from the TRAST measurements, see Figure 1D, with the relative brightness of the P state set to Q = 0 (Eq. S13). Based on this model (and Eqs S5-S13), the individual fluorophore subject to MINFLUX localization can thus occupy the states N (bright trans isomer), P (dark cis isomer), T (dark triplet) and $\dot{R}^{-}$ (dark photo-reduced state) with molecule being in state N at $t = 0$.

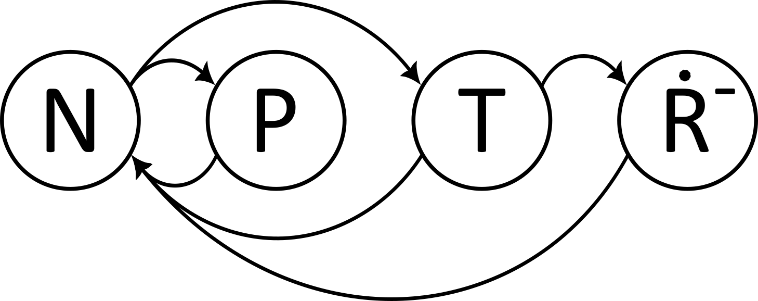

**Figure S4:** Markovian chain model for the fluorophore photophysical states with the possible transitions between the states. The fluorophores undergo fluorescence blinking between the bright trans isomer state (N) and the dark cis isomer (P), triplet (T) and the long-lived redox ($\dot{R}^{-}$) state. The $\dot{R}^{-}$ state is populated through the T state, but relaxes back to the N state, following the model of Figure 1D..

Following the rate matrix for the photophysical transitions, as stated in Eq. S6, the stochastic matrix for this Markovian chain with the rows corresponding to the populations of the four different states of the model, $N, P, T$ and $\dot{R}^{-}$, is given by

$M= \left[ \begin{matrix} 1-\left( k_{iso}^{'}+k_{isc}^{'} \right)\cdot\Delta t & k_{iso}^{'}\cdot\Delta t & k_{isc}^{'}\cdot\Delta t & 0 \\ \left( k_{biso}^{'} \right)\cdot\Delta t & 1-\left( k_{biso}^{'} \right).\Delta t & 0 & 0 \\ k_{T}\cdot\Delta t & 0 & 1-\left( k_{T}+k_{red} \right)\cdot\Delta t & k_{red}\cdot\Delta t \\ k_{ox}\cdot\Delta t & 0 & 0 & 1-k_{ox}\cdot\Delta t \end{matrix} \right]$ (S18)

where the transition rates are directly obtained from the fitting of the TRAST curves (Table S1), and defined as in Section S3 (Eqs. S6-S9). At each $\Delta t$, the row corresponding to the state occupied by the molecule at the previous time step gives the probability of transition to another state or to stay in the same state. The new state is then sampled from a multinomial distribution with the given probabilities. This is repeated for the whole iteration and the effects of changing $k_{01}$ makes the matrix *M* change in each beam position since $k_{iso}^{'}$, $k_{biso}^{'}$and $k_{isc}^{'}$are excitation dependent (Eqs. S6-S9).

Following Eq. S13 (disregarding the overall detection quantum yield of the instrument and any emission from P), the number of photons from each of the beam positions at the end of each TCP iteration is calculated by

$n_{i}=\sum_{j=0}^{j=t_{dwell/\Delta t}} {\Delta tk}_{10}q_{f}S_{1}\left( t \right) with t=i\times t_{dwell}+j\times\Delta t \mathrm{for} i \in[0,1,\ldots K-1]$	(S19)

,with the instantaneous equilibrium excited singlet state S_1_ population of the N state given by

$S_{1}\left( t \right)= \frac{\sigma_{N}\cdot\Phi_{exc}(t)}{\sigma_{N}\cdot\Phi_{exc}(t)+k_{10}^{N}}\cdot N\left( t \right)$	(S20)

Here, $\Phi_{exc}$ denotes the excitation photon flux experienced by the fluorophore within the TCP iteration.

After the calculation of$n_{i}$ for K beam positions the photon counts $\bar{n}= \left\{ n_{0},n_{1},\ldots,n_{K-1} \right\}$ are used for maximum likelihood estimation (MLE) of the fluorophore position (following the same procedure as described in Balzarotti et al, Science, 2017,^11^ supplementary section 1 and 3.1.2). In order to make inferences only on the effects of photophysics on the MINFLUX localizations, microscope and detector specific variability and background is not considered.

In order to estimate the position $\bar{r}_{m}$ of the emitter given the photon count vector $\bar{n}$ , we find the argument which maximizes the likelihood function $\mathcal{L}\left( \bar{r} | \bar{n} \right)=P\left( \bar{n} | N, \bar{r} \right)$which is the conditional probability of measuring the set of photons $\bar{n}$ given the total number of photons N and the position $\bar{r}_{m}$, given by

$\mathcal{L}\left( \bar{r} | \bar{n} \right)=\frac{N!}{n_{0}!\ldots n_{K-1}!}\prod_{i=0}^{K-1} {p_{i}(\bar{r})}^{n_{i}}$	(S21)

Where, for the background free case, $p_{i}\left( \bar{r} \right)$ can be calculated from

$p_{i}\left( \bar{r} \right)= \frac{I_{i}(\bar{r})}{\sum_{0}^{K-1} I_{j}(\bar{r})} with i \in[0, 1, \ldots, K-1]$	(S22)

Since the argument which will maximize the likelihood function will also maximize the log likelihood function, we find the position estimate by finding the argument which will maximize the log likelihood function $l\left( \bar{r} | \bar{n} \right)$ given by

$\ln\mathcal{L}\left( \bar{r} | \bar{n} \right) \propto l\left( \bar{r} | \bar{n} \right)= \sum_{i=0}^{K-1} n_{i}\ln p_{i}$	(S23A)

$\bar{r}_{m}^{MLE}=argmax(l\left( \bar{r} | \bar{n} \right))$	(S23B)

The TCP center is then shifted to the new estimated location of the molecule $\bar{r}_{m}^{MLE}$ for the next iteration where the TCP diameter *L* is reduced while the laser power is ramped up in multiples of the starting power and the procedure is repeated. The power ramp for the different iterations is 1x, 2x, 4x, 6x of the starting power. For reference, at a starting power of 5 µW and with the beam dimensions as stated above the corresponding maximum intensity (in the donut ring) is 3.4 kW/cm^2^. For simplicity, the time taken for the MLE calculation in a real MINFLUX localization between iterations is not considered in the simulation, since the state evolution at that time is strongly depended on the beam resting position, laser power and calculation time which is not universal.

The state evolution of an ensemble of fluorophores at position ${(\bar{r}}_{m})$ instead of a single fluorophore is simulated in a similar manner as for a single fluorophore. However, instead of multinomial sampling with probabilities given by the stochastic matrix in the single molecule case, time evolution of the state vector $\bar{A}= \left[ N, P,T,\dot{R}^{-} \right]^{T}$ is such that

$\frac{\Delta\bar{A}}{\Delta t}=M\left( \Delta t \right). \bar{A}(t)$	(S24)

Where the $M\left( \Delta t \right)$ summarizes the different transition rates from the different states and is updated based on the MINFLUX beam position as in the single molecule case. The matrix is the same as stated in Eq. S6, and is given by

$M= \left[ \begin{matrix} -\left( k_{iso}^{'}+k_{isc}^{'} \right) & \left( k_{biso}^{'} \right) & k_{T} & k_{ox} \\ k_{iso}^{'} & -\left( k_{biso}^{'} \right) & 0 & 0 \\ k_{isc}^{'} & 0 & -\left( k_{T}+k_{red} \right) & 0 \\ 0 & 0 & k_{red} & {-k}_{ox} \end{matrix} \right]$	(S25)

The evolved state vector after any simulation time step, $\Delta t$, is then given by

$\bar{A}_{t}= e^{M\Delta t}\bar{A}_{t-1}$	(S26)

**Section S10:** Comparison of state evolution simulations of individual fluorophores with ensemble state population kinetics

The different implementations for the state evolution for ensemble and single fluorophores were compared to see if the ensemble evolution is recreated if many single molecule evolutions are averaged. The state evolution over ~1ms with a constant $k_{01}=1{\mu s}^{-1}$was simulated for 1000 single DL755 fluorophores (see examples in Figure S5) and then averaged to compare with the ensemble case with the same conditions (Figure S6). The averaged state evolution obtained from the single molecule case, agreed well with the ensemble simulations.

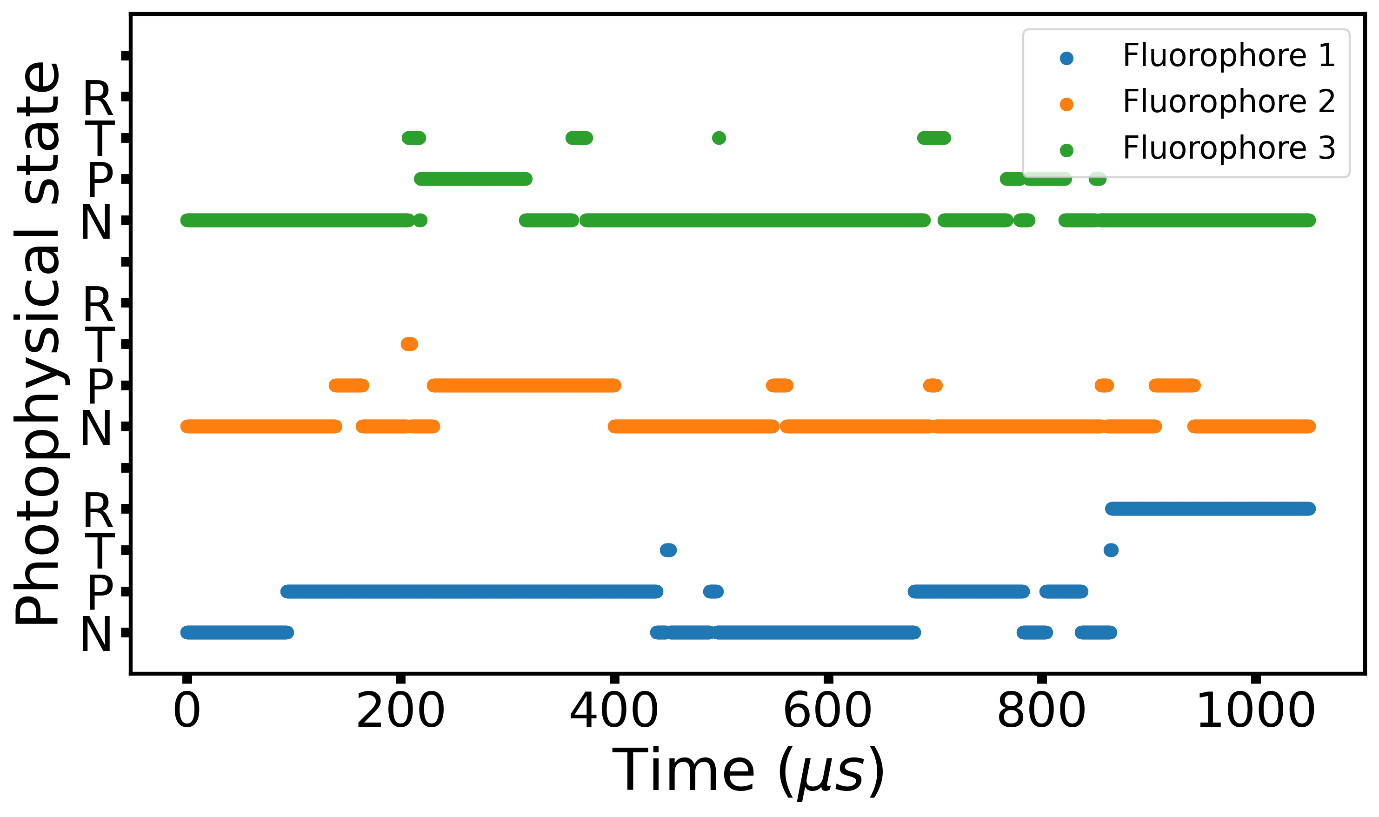

**Figure S5:** Examples of single molecule photophysical state evolutions for three different DL755 fluorophores experiencing constant $k_{01}=1{\mu s}^{-1}$ . The fluorophores undergo fluorescence blinking, generated by transitions between the bright trans isomer state (N) and the dark cis isomer (P), triplet (T) and the long-lived redox (R) state, according to the model in Figure 1D and based on determined rate parameters for DL755 (Table S1)..

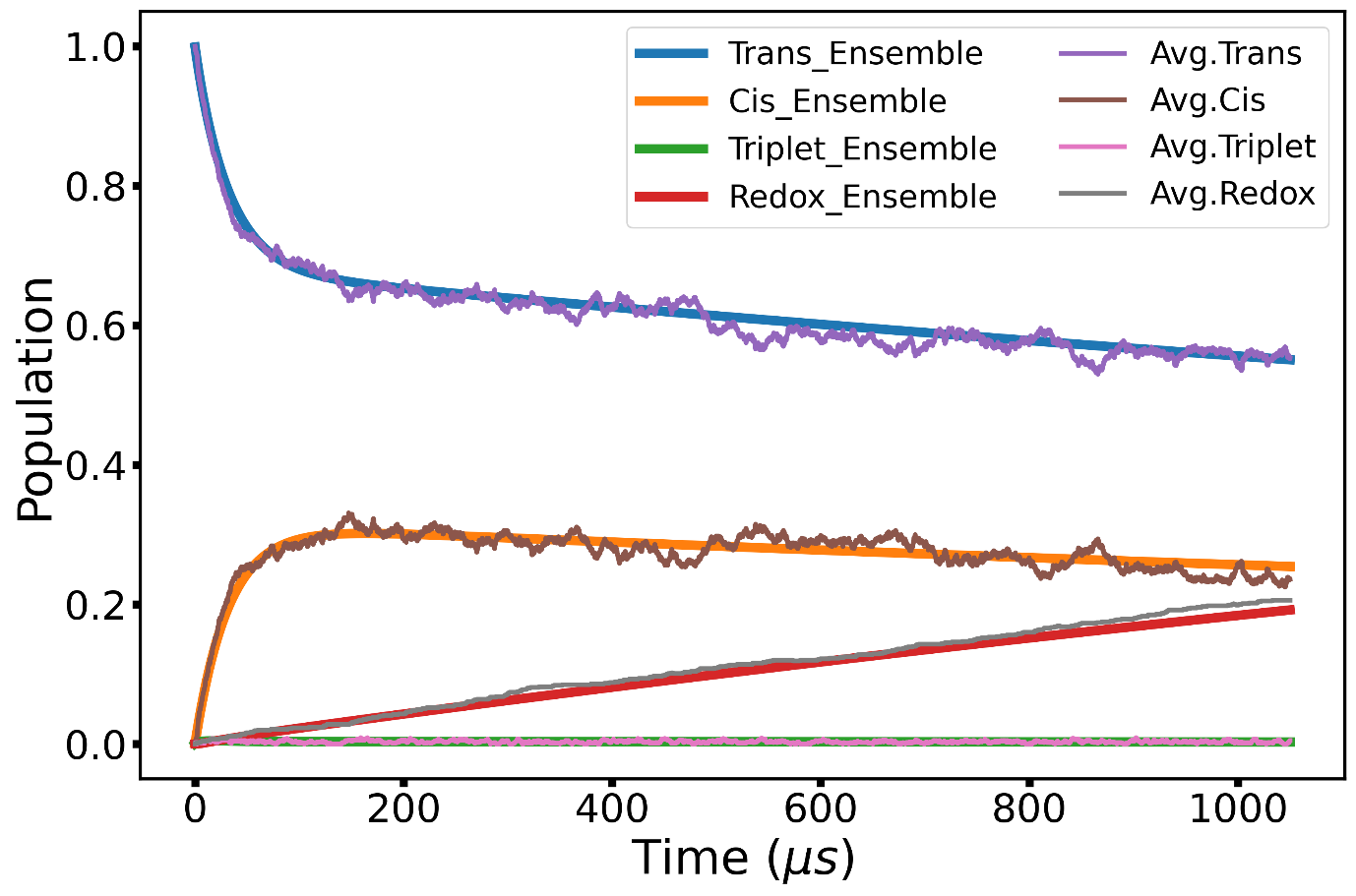

**Figure S6:** Comparison of simulated ensemble and single molecule state evolutions of DL755 at onset of constant excitation with $k_{01}=1{\mu s}^{-1}$. In the case of the single molecule simulations, the population of each state is averaged for 1000 fluorophores.

**Section S11:** Simulated localization errors after individual TCP iterations, for short *t_dwell_* / low $\Phi_{exc}$ and longer *t_dwell_* / higher $\Phi_{exc}$, respectively.

**Figure S7:** Maps of estimated locations (coloured dots) of a fluorophore simulated to be at position (1,1) (golden star). Estimated locations are shown at the end of each iteration of the simulated iterative MINFLUX localizations. Errors in localization and the extent of inaccurate localizations reduced with iterations (DL755 with beam dwell time of 5µs, starting laser power of 10µW and without pattern repeat). The ROI (each box) has the dimensions -100nm to +100nm in the x and y directions.

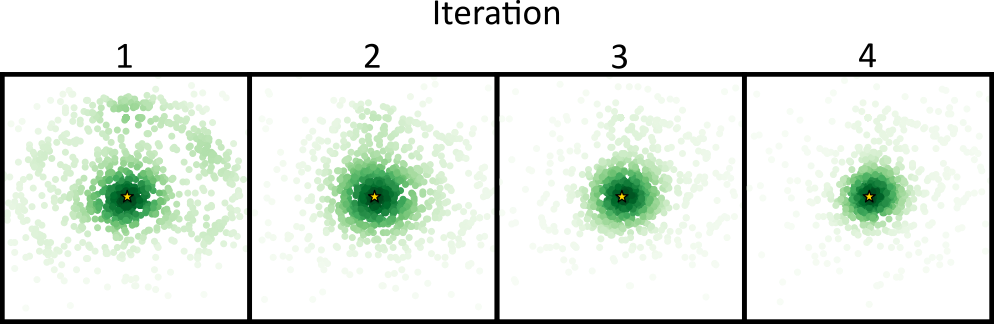

**Figure S8:** Maps of estimated locations (coloured dots) of fluorophore simulated to be at position (1,1) (golden star), simulated as in Figure S4, but now with beam dwell times of 150µs instead of 5µs. In this case, additonal iterations do not improve the large extent. This can be attributed to the fact that in most cases the fluorophore is populated into the long-lived dark redox state during the first iteration and is not recovered until after the simulation time. In some of the simulations, in which the fluorophore came back to the N state, showed improved estimates, to an extent depending on the TCP size and position with respect to the fluorophore position. The ROI (each box) has the dimensions -100nm to +100nm in the x and y directions.

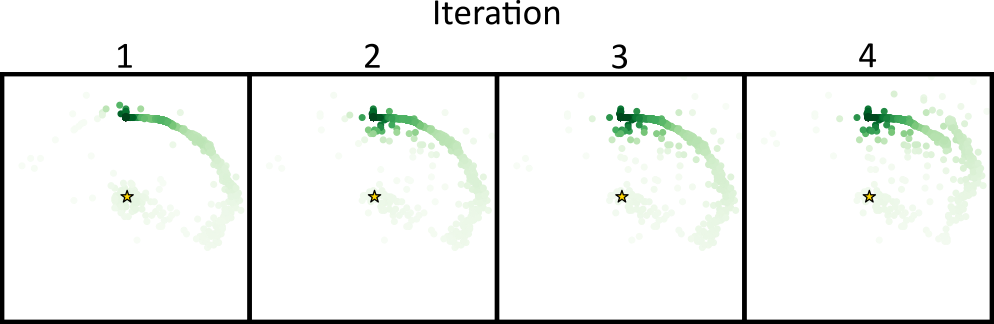

**Section S12:** Simulated localizations for A647

**Figure S9:** Maps of simulated final estimated locations (coloured dots) for a AF647 fluorophore at position (1,1) (golden star) after iterative MINFLUX localization. The figure shows simulated localization errors for different powers, different beam dwell times and without (A) or with (B) pattern repeats (dotted boxes, pattern repeat of 5). With pattern repeats, most of the estimates fall onto the actual position with similar localization errors for different powers and beam dwell times. The ROI (each box) has the dimensions -100nm to +100nm in the x and y directions.

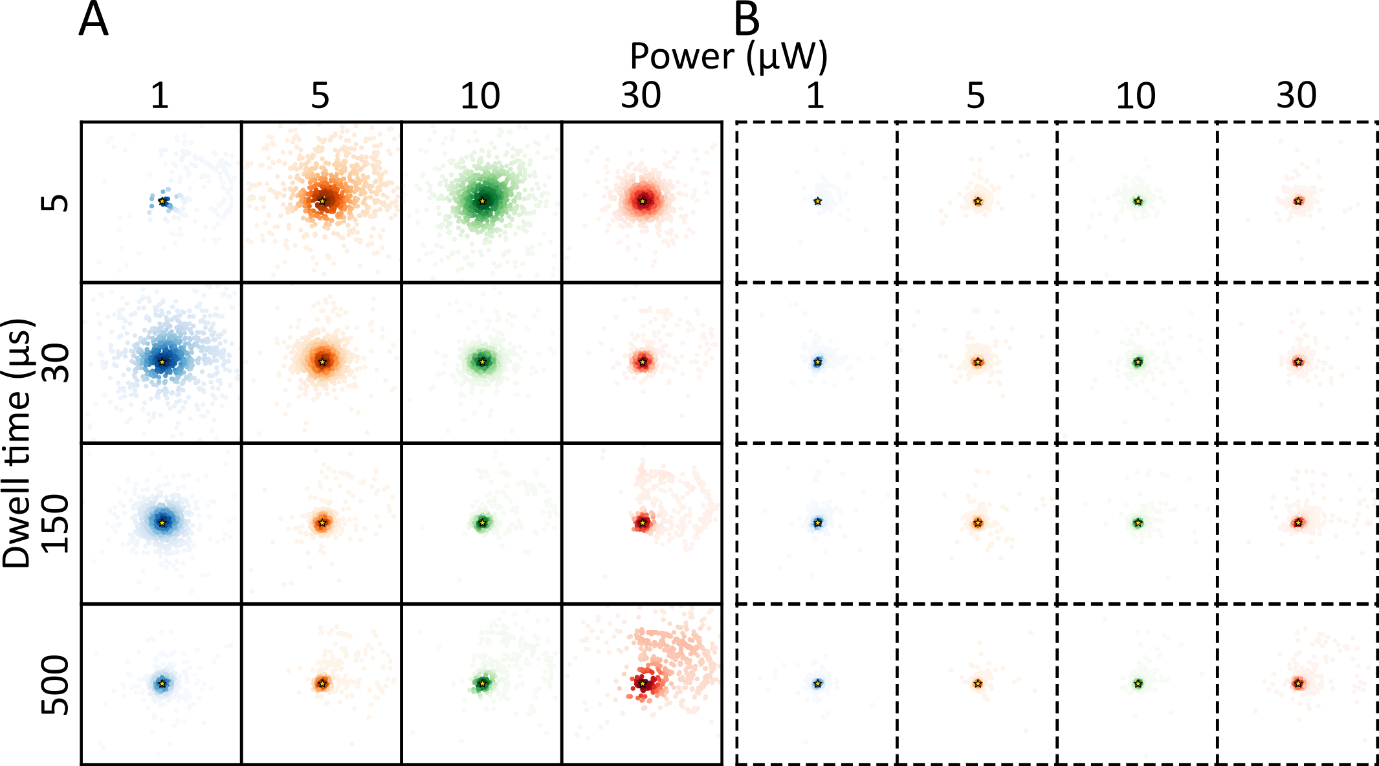

**Section S13:** Simulated localizations for DL755 with a reversed TCP direction and with a different fluorophore location (in relation to the TCP)

**Figure S10:** Maps of simulated final estimated locations (coloured dots) for a DL755 fluorophore at position (1,1) (golden star) after iterative MINFLUX localization. The figure shows the localization error distributions for different powers, different beam dwell times, with different beam directions. The left matrix shows the outcome when the position of the TCP beam is altered in a clockwise direction, and the right matrix for a corresponding counter-clockwise beam direction. ROI (each box) has the dimensions -100nm to +100nm in x and y directions.

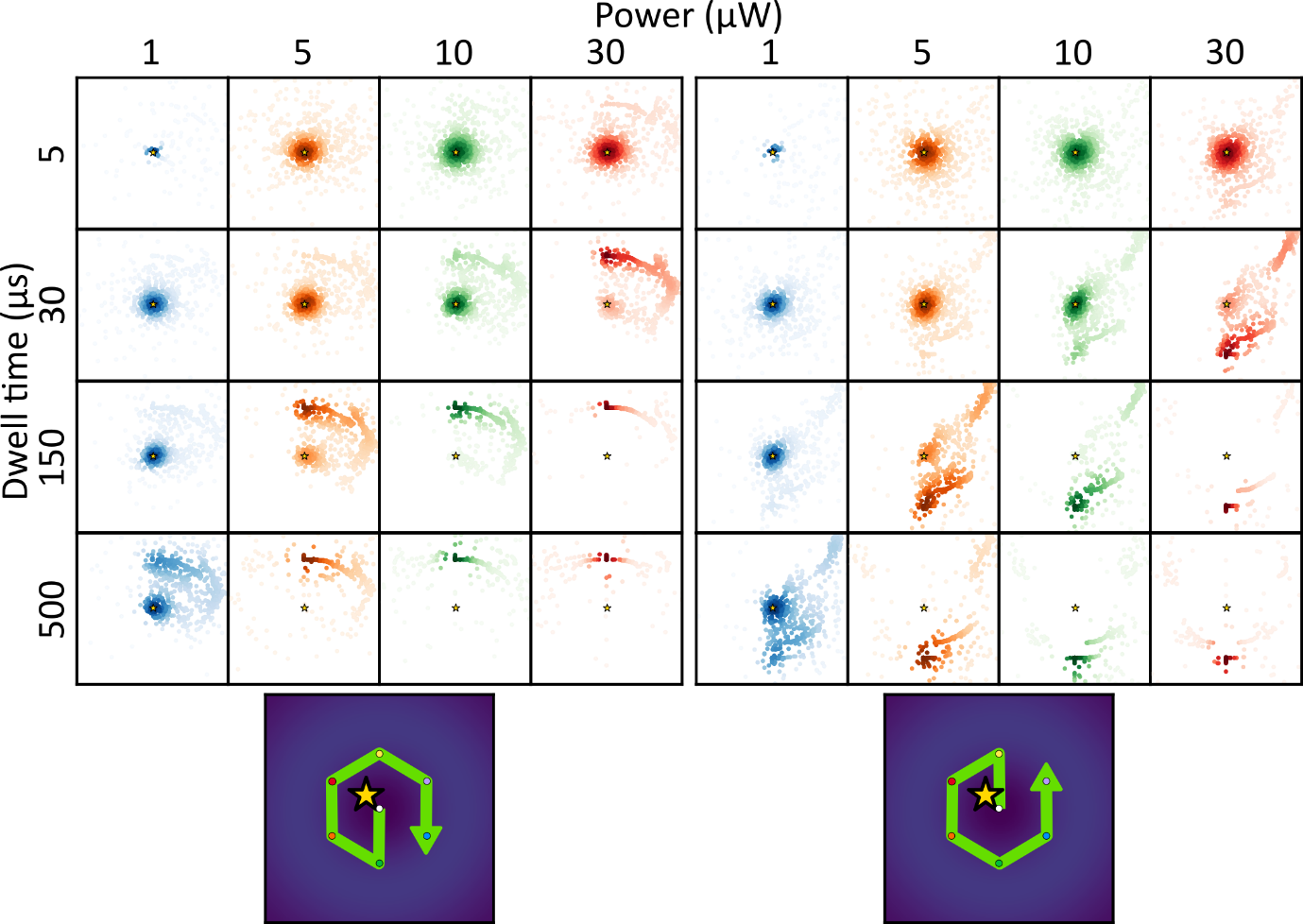

**Figure S11:** Maps of simulated final estimated locations (coloured dots) for a DL755 fluorophore at position (A) (-35,35) (B) (-5,20) (C) (-40,-30) (golden star) after iterative MINFLUX localization. The figure shows the localization error distributions for different powers, different beam dwell times and without (left) or with (right) pattern repeats (dotted boxes, pattern repeat of 5). The figure shows that the spatial distribution of localization errors also strongly depends on the fluorophore position, clearly showing a different pattern of errors with respect to Figure 3, main text. ROI (each box) has the dimensions -100nm to +100nm in the x and y directions.

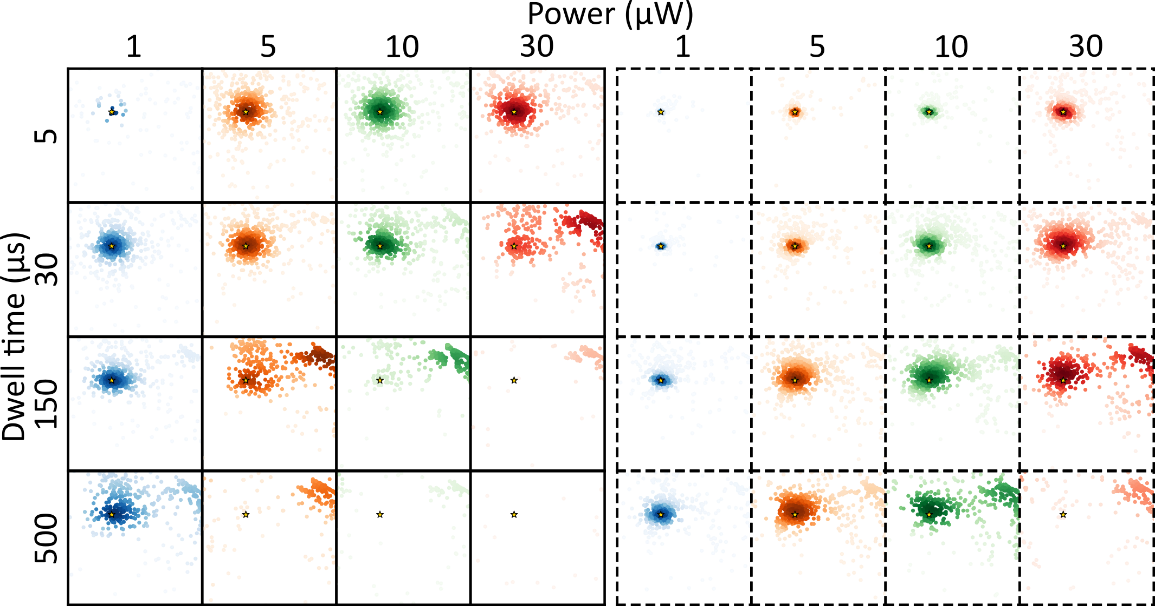

**A**

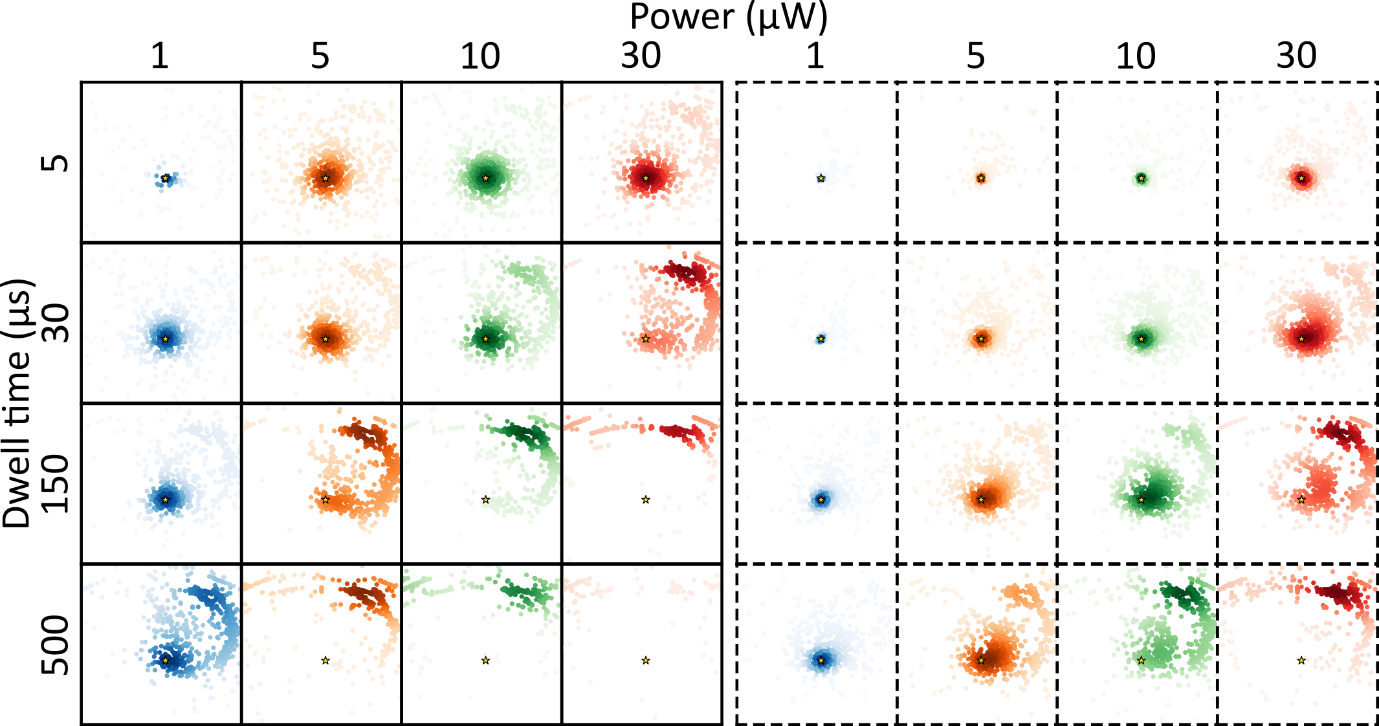

**B**
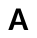

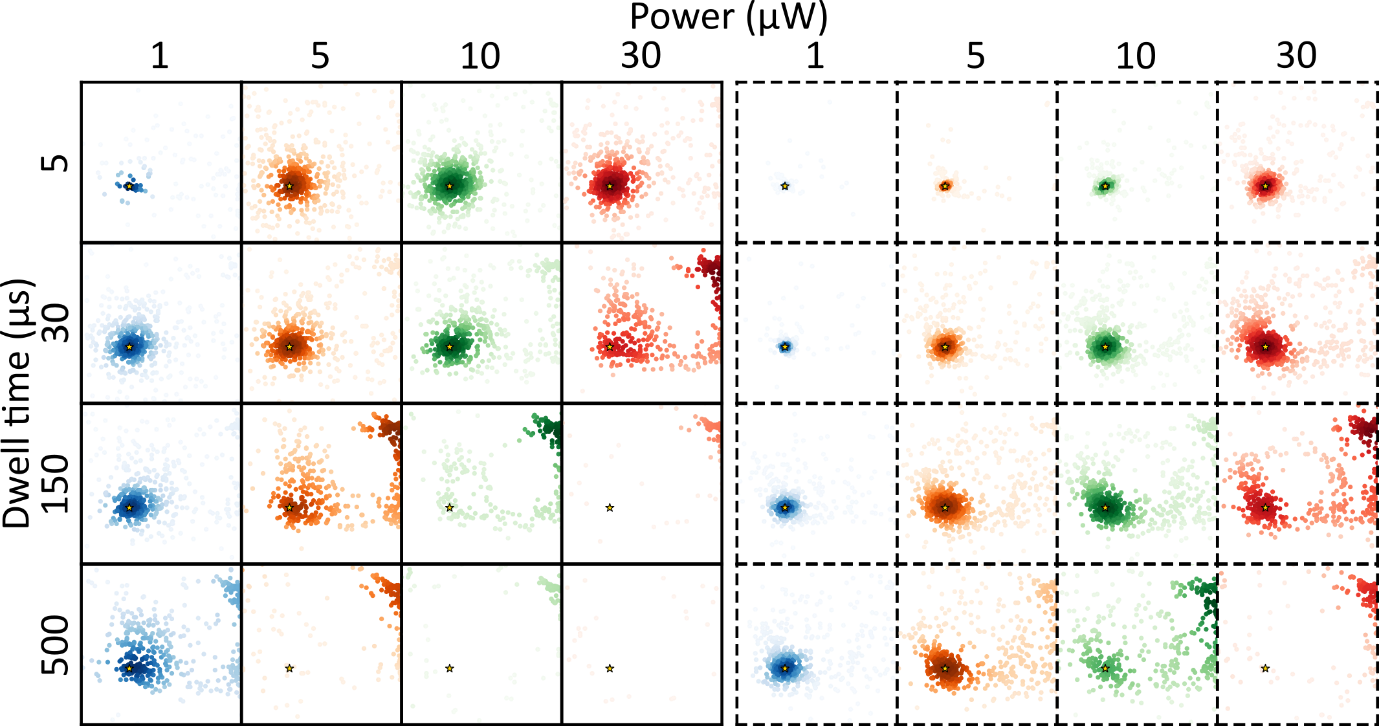

**C**
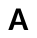

**Section S14:** Simulated localizations for an ensemble of DL755 fluorophores (at the same location)

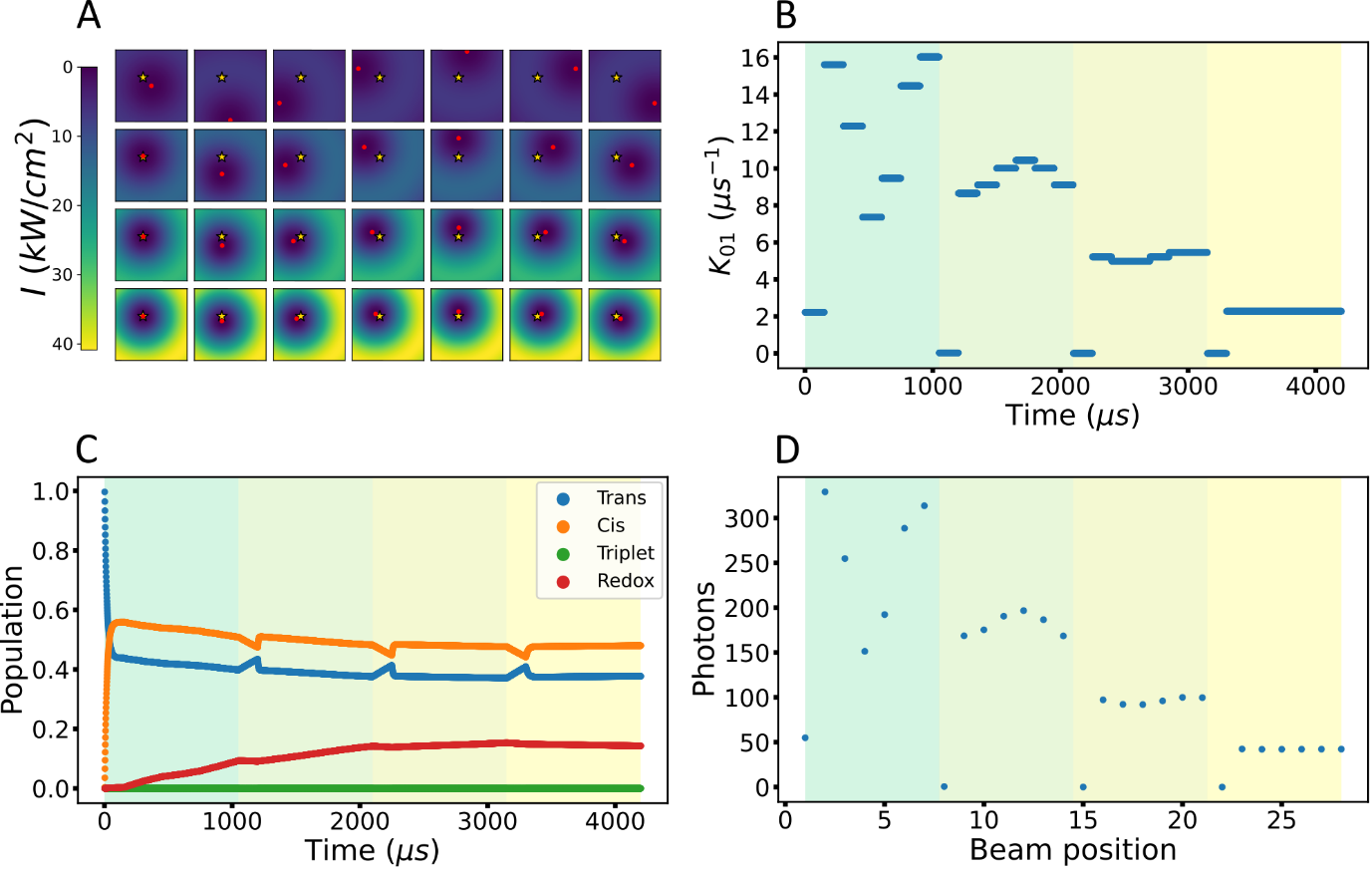

**Figure S12:** Simulation of a MINFLUX localization, based on an ensemble population kinetics, in contrast to the single molecule simulations in Figure 2 (main text). A) Illustration of the fluorophore position (golden star, in this example located at (-35nm,35nm) with respect to the TCP center in the first iteration) and the different beam positions (beam centers marked as red dots) over the different TCP iterations (corresponding to each row), with the excitation intensity increasing and the diameter of the TCP (L) decreasing from one TCP iteration to the next. B) The excitation rate ($k_{01}$) experienced at the fluorophore position over time as the beams are moved in the four TCPs. C) The ensemble photophysical state evolution showing the normalized populations of different states at the corresponding time (N, blue) and the dark cis isomer (P, orange), triplet (T, green) and the dark long-lived redox ($\dot{R}^{-}$, red) state during the time of localization. D) The integrated photons collected from the fluorophores for each of the beam positions shown in figure A. The different coloured regions indicate different iterations in figure B, C and D.

**Figure S13:** Maps of simulated final estimated locations (coloured dots) for an ensemble of DL755 fluorophores at position (1,1) (golden star) after iterative MINFLUX localization. The figure shows simulated localization errors for different powers, different beam dwell times and without (A) or with (B) pattern repeats (dotted boxes, pattern repeat of 5). With pattern repeats, all the estimates fall onto the actual position for different powers and beam dwell times. The ROI (each box) has the dimensions -100nm to +100nm in the x and y directions..

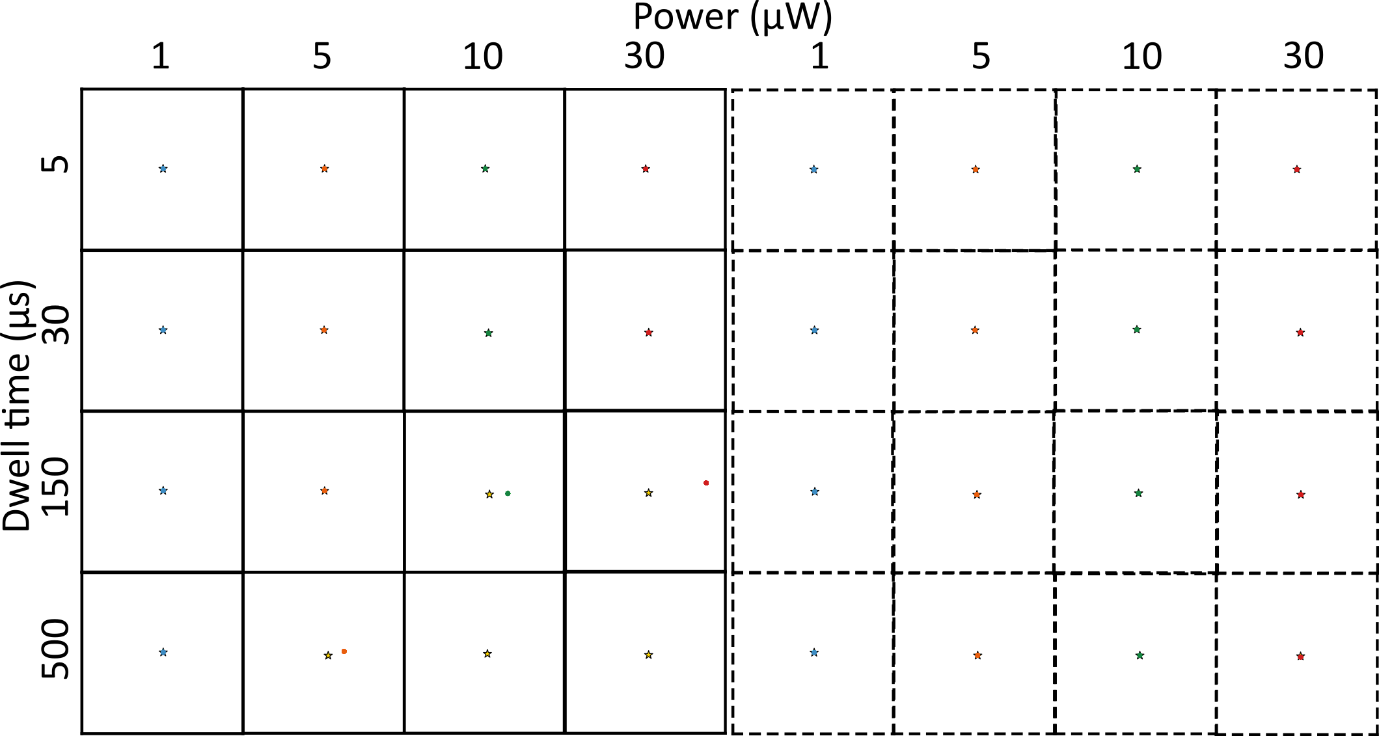

**Figure S14:** The map of simulated estimated locations (coloured dots) of DL755 like fluorophores, with no blinking due to isomerization or photo-reduction, at position (1,1) (golden star) at the end of each iteration of the simulated iterative MINFLUX localization. In the absence of blinking and any background, the simulation can accurately localize the single molecule fluorophores. The ROI (each box) has the dimensions -100nm to +100nm in the x and y directions.

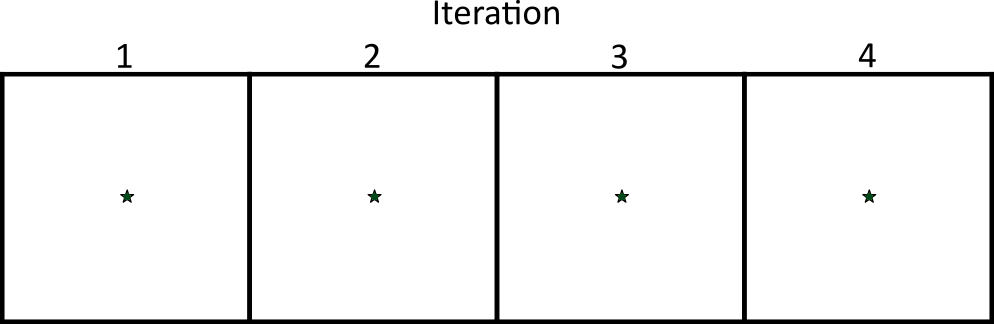

**Section S15:** Simulated localizations for a hypothetical A647 fluorophore with no photo-reduction (isomerization-only)

**Figure S15:** Maps of simulated, final estimated locations (coloured dots) for a AF647-like fluorophore, with no long-lived redox state, at position (1,1) (golden star) after iterative MINFLUX localization. The figure shows the localization errors for different powers, different beam dwell times, and without (A) and with (B) pattern repeats (dotted boxes, pattern repeat of 5). With pattern repeats, most of the estimates are accurately localizing the actual position with similar localization errors between different powers and beam dwell times. The ROI (each box) has the dimensions -100nm to +100nm in the x and y directions.

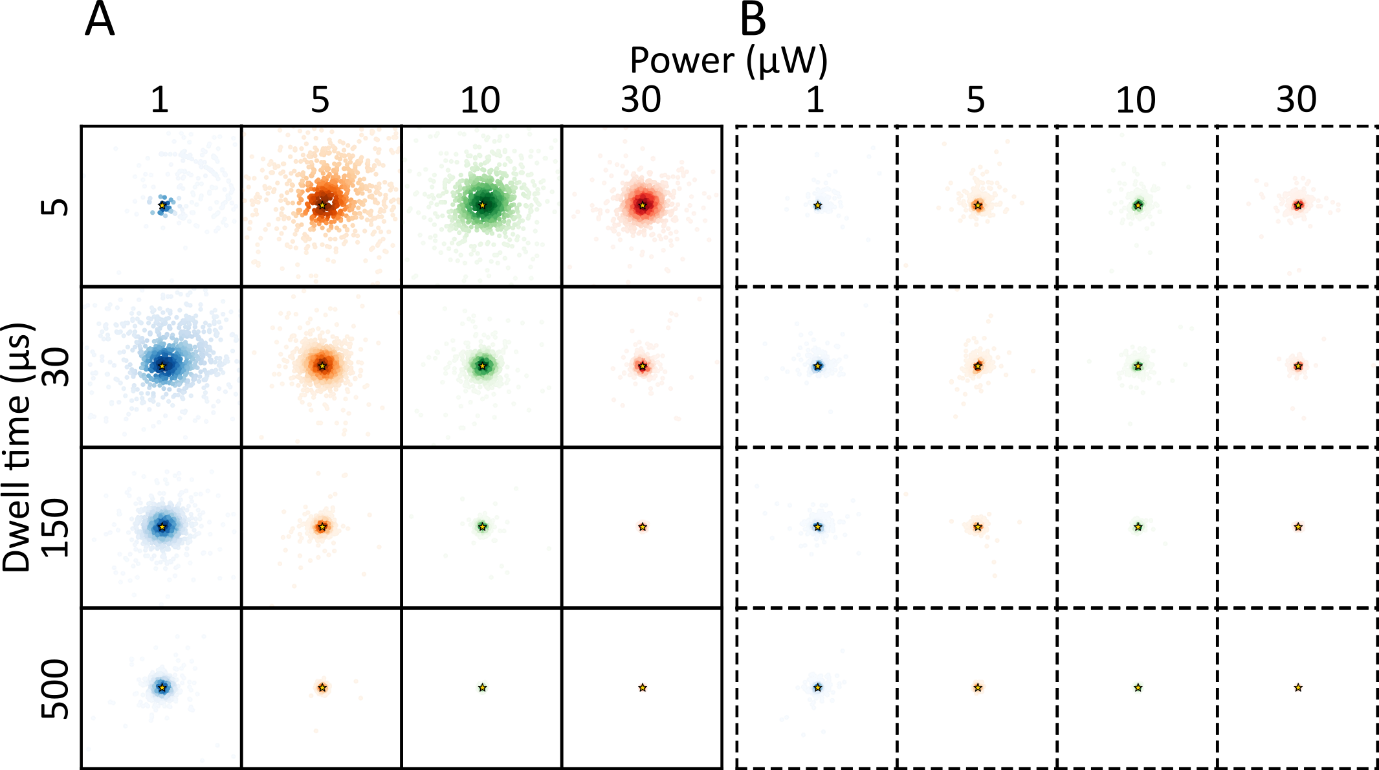

**Section S16:** TRAST curves recorded from DL755 and AF647 in different redox buffers

**Figure S**1**6:** TRAST curves with different excitation intensities applied, from (A) DL755 recorded in TAE buffer with addition of GODCAT and ROXS (1mM MV and 1mM AA), (B) DL755 in Trolox buffer (TAE + GODCAT + 5mM Trolox) and (c) AF647 recorded in Redox buffer (TAE + GODCAT + 1mM MV + 1mM AA )

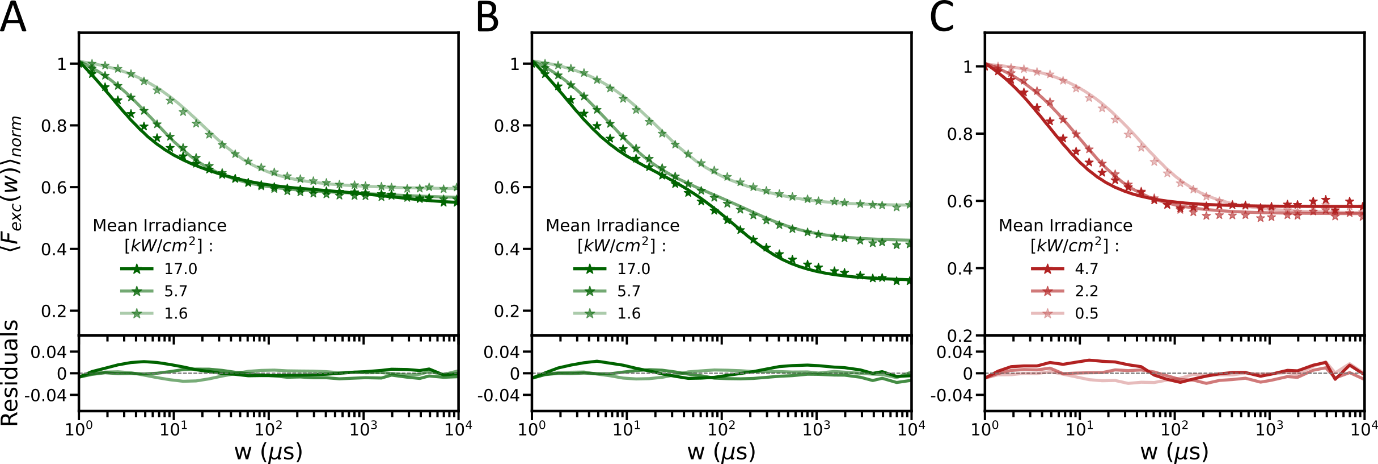

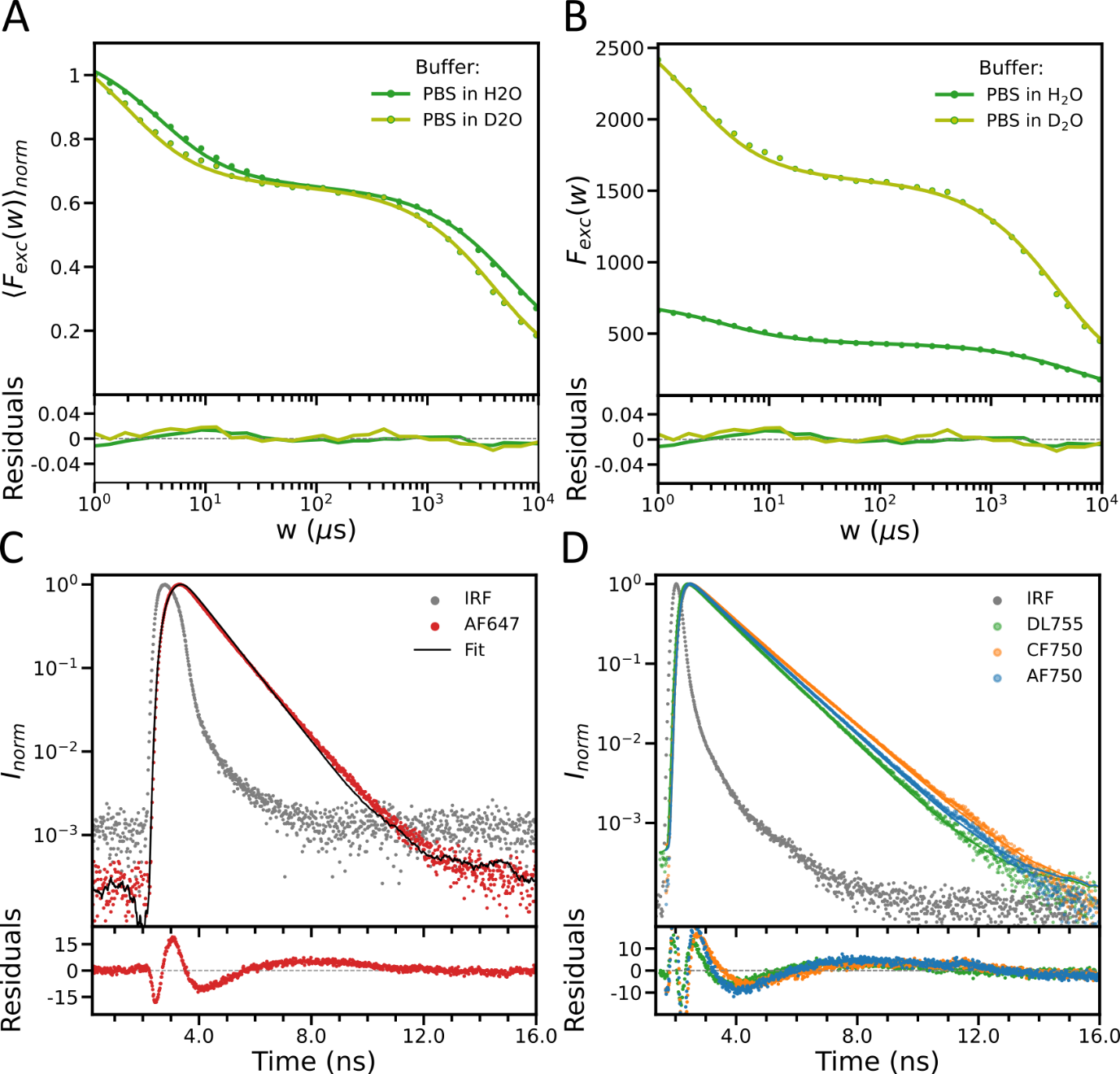
**Section S17:** Effects of heavy water

**Figure S17**: A) Normalized and B) non-normalized TRAST curves from DL755 in PBS using H_2_O or D_2_O as the solvent (Excitation at 750nm, with 4.9 kW/cm). TCSPC measurements from free C) AF647 and D) NIR dyes in D_2_O. The fitted curve for AF647 shows a lifetime of 1.2 ns, compared to 1.07ns in H_2_O (Figure S2A). The lifetimes recorded for DL755, CF750 and AF750 fitted to 1.1, 1.3 and 1.2 ns, respectively, to be compared to 0.47ns, 0.6ns and 0,51ns, as obtained in H_2_O (Figure S2B).

**Section S18: TRAST curves recorded from DL755 in a commercial imaging buffer**

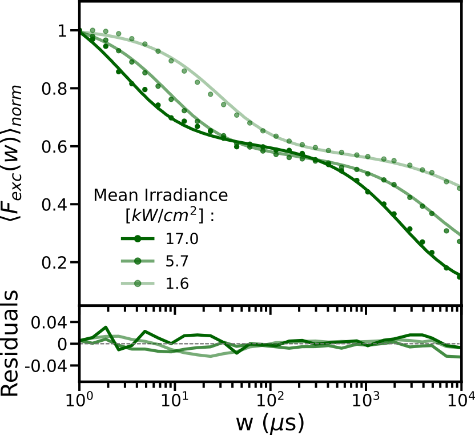

Figure S18: TRAST curve from DL755 conjugated to F1 imager strands recorded in Massive Photonics imaging buffer with different excitation intensities applied.

**Section S19: Nano-ruler MINFLUX images of IR800cw, DY751, AF647**

Figure S19: DNA PAINT MINFLUX image of nanorulers imaged with F1 imager strands conjugated with AF647 recorded in redox buffer (TAE in D20, GODCAT and ROXS).

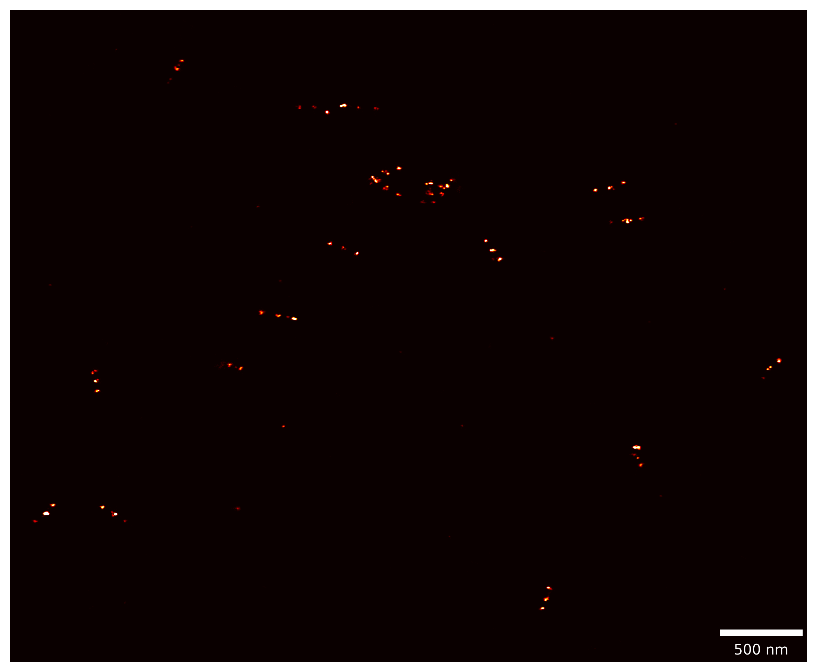

**Section S20: TRAST curve and Nano-ruler MINFLUX images of ATTO700**

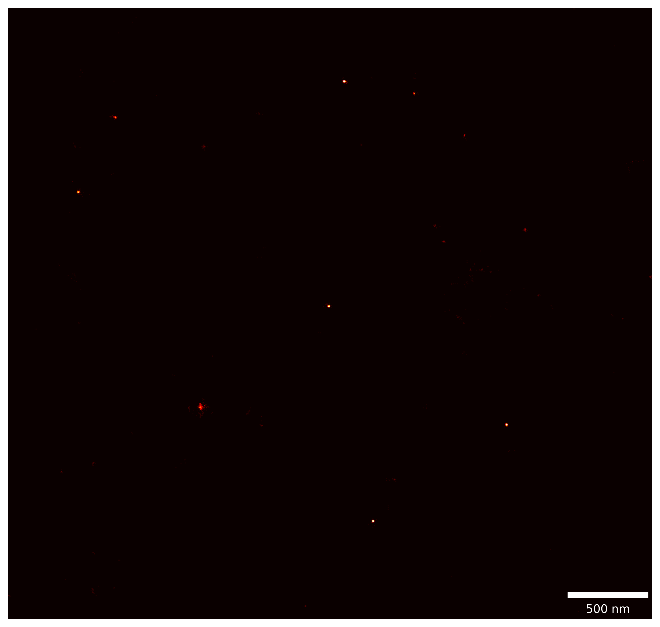

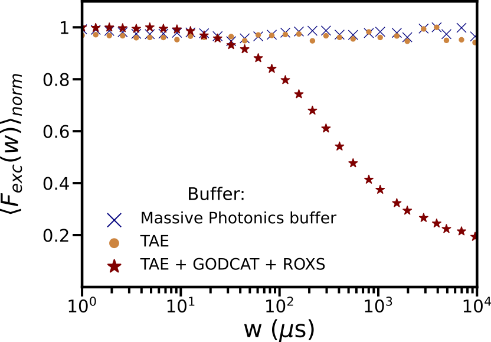

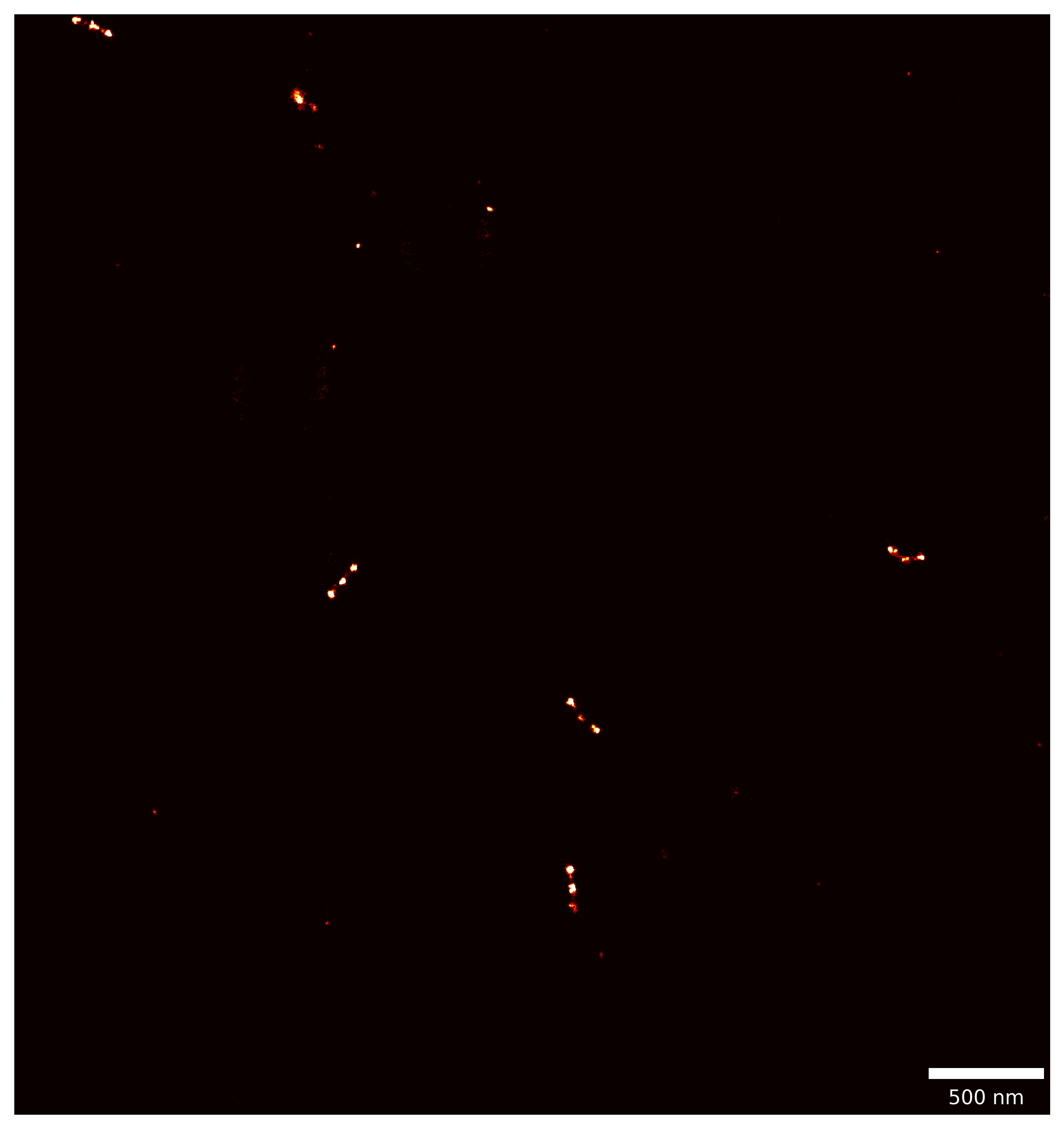

**A**

**B**

**C**

Figure S20: A) TRAST curve from Atto700 conjugated to F1 imager strands recorded in different imaging buffers. DNA PAINT MINFLUX images of nanorulers imaged with F1 imager strands conjugated with Atto700 dye recorded in B) Commercial Massive Photonics DNA-PAINT buffer C) redox buffer (TAE in D20, GODCAT and ROXS).

**Section S21: TRAST curve recorded from DL755 in a deuterated ROXS redox buffer, placed in a non-sealed well chamber open to air exchange.**

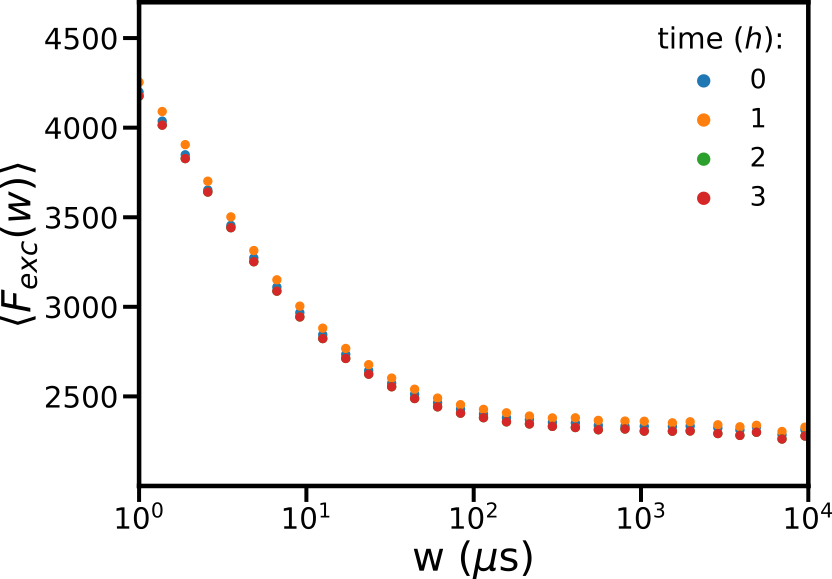

Figure S21: Non-normalized TRAST curves from DL755 in a deuterated ROXS redox buffer, placed in a non-sealed well chamber, open to air exchange, and recorded at 0 to 3 hours after application of the sample in the well chamber. No effects over time were observed, indicating that neither photophysical transitions, nor the absolute brightness of the dye was affected.

**Section S22: MINFLUX sequence used in simulations and imaging experiments**

| **Imaging 2D**  **Seq** | **TCP parameter L** | **Minimum photon count** | **Dwell time (ms)** | **Pattern repeat** | **CFR limit** | **Laser power factor** |
| --- | --- | --- | --- | --- | --- | --- |
| **Pre-localization** | **288** | **160** | **≥1** | **1** | **2** | **1** |
| **Iteration 1** | **288** | **150** | **≥1** | **5** | **Off** | **1** |
| **Iteration 2** | **151** | **100** | **≥1** | **5** | **0.8** | **2** |
| **Iteration 3** | **76** | **100** | **≥1** | **5** | **0.8** | **4** |
| **Iteration 4** | **40** | **150** | **≥1** | **5** | **2** | **6** |
